# Lowering levels of reelin in entorhinal cortex layer II-neurons results in lowered levels of intracellular amyloid-β

**DOI:** 10.1101/2022.01.28.478143

**Authors:** Asgeir Kobro-Flatmoen, Claudia Battistin, Rajeevkumar Nair Raveendran, Christiana Bjorkli, Belma Skender, Cliff Kentros, Gunnar Gouras, Menno P. Witter

## Abstract

Projection neurons in the anterolateral part of entorhinal cortex layer II (alEC LII) are the predominant cortical site for hyperphosphorylation of tau (p-tau) and formation of neurofibrillary tangles (NFTs) in brains of subjects with early-stage Alzheimer’s Disease (AD). A majority of alEC LII-neurons are unique among cortical excitatory neurons by expressing the protein reelin (Re+). In AD patients, and a rat model for AD overexpression mutated human APP, these Re+ excitatory projection-neurons are prone to accumulate intracellular amyloid-β (iAβ). Biochemical pathways that involve reelin-signaling regulate levels of p-tau, and iAβ has been shown to impair such reelin-signaling. We therefore used the rat model and set out to assess whether accumulation of iAβ in Re+ alEC LII projection neurons relates to the fact that these neurons express reelin. Here we show that in Re+ alEC LII-neurons, reelin and iAβ42 engage in a direct protein-protein interaction, and that microRNA-mediated lowering of reelin-levels in these neurons leads to a concomitant reduction of non-fibrillar iAβ ranging across three levels of aggregation. Our experiments are carried out several months before plaque pathology emerges in the rat model, and the reduction of iAβ occurs without any substantial associated changes in human APP-levels. We propose a model positioning reelin in a sequence of changes in functional pathways in Re+ alEC LII-neurons, explaining the region and neuron-specific initiation of AD pathology.

**Significance:** Anterolateral entorhinal cortex layer II (EC LII) neurons are the predominant cortical site for hyperphosphorylation of tau (p-tau) and formation of neurofibrillary tangles (NFTs) in brains of subjects with early-stage Alzheimer’s disease (AD). The same neurons are prone to very early accumulation of non-fibrillary forms of amyloid-β in the context of AD, and are unique among cortical excitatory neurons by expressing the protein reelin. We show that in such alEC LII-neurons, reelin and iAβ42 engage in a direct protein-protein interaction, and that selectively lowering levels of reelin leads to a concomitant reduction of non-fibrillar Aβ. We propose a model positioning reelin in a sequence of changes in functional pathways in reelin-expressing EC LII neurons, explaining the region and neuron specific initiation of AD.

## Introduction

Amyloid-β (Aβ) is an aggregation-prone peptide of 37-43 amino acids in length that is normally present in low levels throughout the brain. The peptide is the result of a sequential cleavage of the amyloid precursor protein (APP) that normally occurs at a low rate. Subtle differences in this cleavage sequence result in variations in the length of Aβ(1). Recent studies on large human cohorts support the concept that the common, sporadic form of Alzheimer’s disease (AD) starts with an increase of Aβ in the brain, and that this leads to pathological hyperphosphorylation of tau (p-tau)(2, 3), eventually resulting in formation of neurofibrillary tangles (NFTs). This concept is substantiated by a number of findings in patients. First, familial, genetically determined forms of the disease are driven by increased levels of total Aβ, or an increase of the more aggregation-prone Aβ42 peptide relative to Aβ40(4). Second, Down syndrome most often leads to full blown AD-pathology by middle age(5) that is due to an added copy of the part of chromosome 21 containing the *APP-gene*(6), which results in overexpression of APP and a consequent overproduction of Aβ. Third, a genetic variant in humans, resulting in less Aβ, protects against AD(7).

Within the AD-research field, the role for Aβ during prodromal AD is most often ascribed to that of its fibrillar-and deposited extracellular forms, colloquially referred to as Aβ-plaques(1, 8, 9). In broad strokes, the notion is that the emerging Aβ-plaques damage nearby synapses in a way that induces p-tau. Then, as the degree of hyperphosphorylation reaches beyond a currently unknown threshold, p-tau detaches from and disrupts the integrity of microtubules. A consequent degeneration of the affected axon terminals follows, and the now detached p-tau translocates or misallocates to somatodendritic compartments, where it builds up and fibrillates into NFTs(1). But such a role for Aβ-plaques is difficult to reconcile with the evidence. Aβ-plaques arise first in the neocortex, typically in the medial orbitofrontal and posterior cingulate areas, quickly followed by the precuneus(10). The cortical onset of NFTs is predominantly restricted to the anterolateral portion of the entorhinal cortex (alEC), in neurons residing superficially in layer II (LII)(11-15). The presumed relationship between Aβ-plaques and NFT-onset implies that alEC layer II-neurons have projections forming synapses with neurons in the above-mentioned cortical regions. However, findings in rodents(16, 17) and primates(18-23) show that projections from EC to the medial orbitofrontal cortex, posterior cingulate cortex, and the precuneus, originate predominantly in layer V of EC and only very sparsely or not at all in LII (17, 24-26).

Converging lines of evidence from studies of the human brain using live imaging(10, 27), immunohistochemistry(28-31) and biochemistry(32-37), supported by experimental results from rodent models(38-41) and cell models(38, 42), point to a role for Aβ in non-fibrillated forms in the onset of AD. Such a role likely involves interference with cellular homeostatic and metabolic processes(43) that are neuron-type specific(44), implying that the effect of Aβ upon a given population of neurons is not generalizable to all neurons. Using a transgenic mutated human APP-based rat model(45), we uncovered evidence in support of this notion in that cortical neurons start to accumulate intracellular Aβ (iAβ) up to eight months before the model develops the first Aβ-plaques. In EC, this early accumulation of iAβ is restricted to a subpopulation of LII-neurons, and these constitute projection-neurons that express the glycoprotein reelin. We furthermore analyzed human subjects with early NFT-pathology and this provided corroborating data (Braak stages I-V)(39). This raised our interest in the role of reelin in the onset of AD.

The expression of reelin in projection-neurons is atypical for cortex, where reelin is mainly expressed by scattered interneurons(46), along with occasional layer V pyramidal neurons plus superficial pyramidal neurons of the piriform cortex(47). The reelin-expressing (Re+) EC LII-neurons originate the main afferents of the hippocampal dentate gyrus (DG) and fields CA3/2(48, 49), meaning these EC LII-neurons occupy a central position vis a vis the role of entorhinal-hippocampal interactions in declarative memory formation(50). Multiple lines of evidence further relate reelin to memory formation(51-53). Reelin binds to the apolipoprotein E receptor 2 (ApoER2) and to the very low density lipoprotein-receptor (VLDLR), triggering a signaling-cascade that acts on glutamatergic receptors to enhance glutamatergic transmission(52). Of these two receptors, ApoER2 is considered the main receptor in the brain(54). The potency of this reelin-signaling cascade has become evident in experiments where single-injections of reelin into the ventricles of mice resulted in increased hippocampal dendritic spine density, long-term potentiation (LTP), and enhancement of memory-dependent task-performance(51). Corollary findings showed that reduced levels of reelin associate with impaired memory-dependent task-performance(55).

A series of studies suggest that Aβ might directly affect reelin. Reelin is cleaved to result in a subset of fragments(56), of which the 180-kDa fragment was found substantially increased upon examination of frontal cortices of AD-subjects(57). An even greater increase of the 180-kDa fragment was found in middle aged subjects with Down syndrome (58). A possible interaction between reelin and Aβ was first shown in experiments where synthetic Aβ42 was added to cultured SH-SY5Y neuroblastoma cells, causing a dose-dependent increase of the 180-kDa reelin-fragment in the absence of changes to reelin messenger RNA levels(58). Further studies revealed that reelin extracted from cortices of AD-subjects tends to be of higher molecular mass than the active, signaling-competent form, indicating an interaction of reelin with another protein(59). That reelin was associated with oligomeric Aβ is likely in view of reports that both proteins co-immunoprecipitate in cortical extracts from AD patients(60).

Alongside enhancement of glutamatergic transmission as outlined above, the binding of reelin to ApoeR2 and VLDLR also triggers a parallel signaling cascade that culminates with potently inhibiting the activity of glycogen synthase kinase 3β (GSK3β)(61-63). As GSK3β is required to induce long-term depression(64), the inhibitory effect of reelin upon GSK3β may help to facilitate LTP. Another crucial role of GSK3β is that it acts as one of the main kinases that phosphorylates tau(65). Of direct relevance to this are studies showing that an interaction with Aβ impairs the signaling-capacity of reelin, which impairs the inhibitory control exerted by reelin-signaling upon the activity of GSK3β. This leads to constitutive tau-phosphorylation by GSK3β, and, importantly, this whole sequence of events is known to result in p-tau(59-63).

This body of evidence led us to design a comprehensive set of experiments to test whether *the selective vulnerability of Re+ alEC LII-neurons to accumulate iAβ is linked to their expression of reelin*. In the present study we show that in Re+ alEC LII-neurons, reelin and iAβ42 engage in a direct protein-protein interaction, and that selectively lowering levels of reelin expression leads to a concomitant reduction of several forms of iAβ. The reduction of iAβ occurs without any substantial associated changes in human APP (hAPP)-levels and prior to any plaque pathology in the rat model.

Our findings have important implications for understanding the onset of AD. Increased levels of Aβ occur very early in the initiating phase of the disease in Re+ alEC LII-neurons(39), and the resulting interaction between reelin and Aβ very likely impairs the signaling-capacity of reelin, which, as outlined above, can result in p-tau. It is furthermore safe to assume that the Re+ alEC LII-neurons, which project to the hippocampal formation, are the first cortical neurons to die in AD(66, 67). We propose a model positioning reelin in the sequence of changes in functional pathways in Re+ alEC LII-neurons resulting in the initiation of AD pathology.

## Methods

### Experimental design

We used the McGill-R-Thy1-APP homozygous transgenic rat model, which carries a transgene containing human APP751 with the Swedish double and Indiana mutations expressed under the murine Thy1.2 promoter(40). Animals were bred at the Kavli Institute for Systems Neuroscience at the Norwegian University of Science and Technology. All protocols are approved by the Norwegian Animal Research Authority, complying with the European Convention for the Protection of Vertebrate Animals used for Experimental and Other Scientific Purposes.

Animals were genotyped using quantitative PCR (qPCR), as previously described (68). We used genomic DNA isolated from ear tissue with a High Pure PCR Template Preparation Kit (Roche Diagnostics, Basel, Switzerland). The transgene was detected using RT2 qPCR Primer Assays from Qiagen (Venlo, Netherlands), and as a normalization gene we used (GAPDH or beta-actin), both with FastStart Universal SYBR Green Master (Roche Diagnostics) on an Applied Biosystems StepOnePlus real-time PCR system (Life Technologies Ltd., Thermo Fisher Scientific, Waltham, MA, USA). From the qPCR, ΔΔCT values were calculated with a known homozygous sample as reference (69).

### Adeno Associated Viral (AAV)-vector design

The backbone construct of pAAV-CMV-βglobin-intron-MCS-WPRE-hGH PolyA was made using the DNA sequence of Woodchuck hepatitis virus Post-transcriptional Regulatory Element (WPRE), synthesized and cloned after the multicloning site in pAAV-MCS (Agilent USA, #240071). We then made the control construct pAAV-CMV-βglobin-intron-EGFP-WPRE-hGH PolyA by cloning the EGFP gene sequence between EcoR1 and BamHI restriction sites in the backbone (*the control vector*). Multiple candidate pre micro RNA (miRNA) sequences for knocking-down expression of the *reelin gene* were designed (miRNA-Re) using the RNAi designer tool (Thermo Fisher Scientific, USA) and different pAAV plasmid constructs to express miRNA-Re were created. The pre-microRNA constructs were engineered with endogenous murine miR-155 flanking sequences and were integrated after the EGFP sequence using BamHI and HindIII sites in the pAAV-CMV-βglobin-intron-EGFP-WPRE-hGH PolyA backbone. The positive clones were confirmed by restriction digestion analyses and subsequently by DNA sequencing. Two pAAV plasmid constructs, pAAV-CMV-βglobin-intron-EGFP-miR-RE1 and pAAV-CMV-βglobin-intron-EGFP-miR-RE4 targeting different mouse reelin sequences, that exhibited efficient reelin knockdown in a heterologous cell-culture system based in vitro knockdown assay were chosen for preparing the AAV viral vectors for further in vivo studies (*the experimental vector*).

Endotoxin free plasmid maxipreps (#12663, Qiagen) were made for AAV preparations. The viral vectors were packaged in AAV serotype 2/1 capsids (a mosaic of capsid 1 and 2) and purified using Heparin column affinity purification(70). Briefly, the day before transfection, 7x 106 AAV 293 cells (#CVCL_6871, Agilent, USA) were seeded in DMEM (# 41965062, Thermo Fisher Scientific) containing 10% fetal bovine serum (#16000-044, Thermo Fischer Scientific) and penicillin/streptomycin antibiotics (#15140122, Thermo Fisher Scientific) into 150 mm cell culture plates. Calcium chloride mediated co-transfection was done with 22.5 µg pAAV-containing the transgene, 22.5 µg pHelper (#240071, Agilent, USA), 11.3 µg pRC (#240071, Agilent, USA), and 11.3 µg pXR1 (NGVB, IU, USA) capsid plasmids. The medium was replaced with fresh 10% FBS containing DMEM, 7 hours post-transfection. The cells were scrapped out after 72 hours, then centrifuged at 200xg and the cell pellet was subjected to lysis using 150 mM NaCl-20 mM Tris pH 8.0 buffer containing 10% sodium deoxycholate. The lysate was then treated with Benzonase nuclease HC (#71206-3, Millipore) for 45 minutes at 37°C. Benzonase treated lysate was centrifuged at 3000xg for 15 min and the clear supernatant was then subjected to HiTrap® Heparin High Performance (#17-0406-01, GE) affinity column chromatography using a peristaltic pump (McClure C JOVE 2011). The elute from the Heparin column was then concentrated using Amicon Ultra centrifugal filters (#Z648043, Millipore). The titer of viral stock was determined as approximately 1011 infectious particles/mL.

### Stereotaxic injections

We used 24 animals (10 males, 14 females) for stereotaxic injections, of which we injected five bilaterally and 19 unilaterally with an *experimental vector* (*miRNA-Re EGFP virus*) vs a *control vector* (*EGFP-only virus*). We injected all animals a few days past 1 month of age, when their weight was ∼100 grams (SI Appendix Table 1). We anesthetized animals using 5 % isoflurane gas (Abbott Lab., Cat# 05260-05) in an induction chamber and then immediately transferred the animals to a stereotaxic frame with a mask providing a flow of 1,5-2% isoflurane gas for the full length of the surgery.

To target alEC layer II with high precision, we aligned our coordinates to the brains’ sagittal and transverse sinuses, as their attachment to the brain constitute the most stable anchoring points. Specifically, we horizontally leveled the skull in the stereotaxic frame by lowering the tip of the capillary on top of bregma and then adjusting until lambda was at the exact same horizontal position vis a vis the tip of the capillary. We then aligned the capillary tip to the sagittal sinus at the mid-point between bregma and lambda, and from here moved 3.30 mm laterally, and then caudally until crossing the point where the transverse sinus passes, where the capillary tip was again aligned. From this point the tip was moved 4,60 mm rostrally, and a further 3,60 mm laterally. At this coordinate a small hole was drilled and the capillary tip was lowered to the surface of the brain, from which point the tip was lowered 4,50 mm to hit alEC layer II. We then waited for 5 minutes to alleviate any deflections in the brain tissue, before injecting.

Bilateral injections consisted of one injection of the *experimental vector* into either the left or the right alEC, and one injection of the *control vector* into the contralateral alEC. For the unilateral injections, we injected either the experimental or control vector alternately into the left vs right alEC per animal. The volume of the injections ranged from 300-900nl, and was delivered using a micropump (Drummond Nanoject III, Cat# DRUM3-000-207) with a back filled ultrathin glass capillary at 30 nl/min. After injecting, we waited 5 minutes to allow good uptake before slowly retracting the capillary. We then waited for ∼2 months (SI Appendix Table 1) before processing the animals.

### Brain extraction, tissue processing and immunohistochemistry (IHC)

Animals ranged in age from P54 to P78 at the time of euthanasia (SI Appendix Table 1). We anesthetized animals using isoflurane gas in an induction chamber, followed by intraperitoneal injection of pentobarbital (Norwegian Pharmacy Association, Cat# 306498). Subsequently, we transcardially perfused animals with a Ringer’s Solution (145 mM NaCl, VWR Int. LLC, Cat# 27800.291; 3.35 mM KCl, Millipore, Cat# 1.04936.1000; 2.38 mM NaHCO3, Millipore, Cat# 1.06329.1000), oxygenated to pH ∼6,9, followed by circulation of 4% freshly depolymerized paraformaldehyde (Millipore, Cat# 1.04005.1000) in phosphate buffer (PB: purified de-ionized water with di-sodium hydrogen phosphate dihydrate, Millipore, Cat# 1.37036.500, mixed with sodium di-hydrogen phosphate monohydrate, Millipore, Cat# 1.06346.1000, at 125 mM, pH 7.6; note that this applies to all uses of PB) for 2–3 min. We extracted the brains and post fixed each in the same fixative overnight and then placed them in a freeze protective solution containing 2% DMSO and 20% glycerine (PB with dimethyl sulfoxide, VWR Int. LLC, Cat# 23486.297, and glycerine, VWR, Cat# 24387.292) in a refrigerator until sectioning. Brains were sectioned at 40 μm (for double injected) or 30 μm (for single injected) in the coronal plane with a freezing microtome (Microm HM430, Thermo Fisher Scientific). We collected six (for double injected) or 8 (for single injected) series of equally spaced sections, randomly assigned one series to each IHC experiment, and did all incubations on free-floating sections.

Prior to IHC, all tissue was subject to Heat Induced Antigen Retrieval, by immersion in PB at 60 °C for 2 hours. We blocked tissue with 5% goat serum (2% for 1D1; Abcam, Cat# AB7481) in PB for 1 h. Subsequently, we did double-IHC labeling with primary antibodies (PA) in PB solution containing either 0.2% Triton X-100 (Millipore, Cat# 1.08603.1000) or 0.4% Saponin (VWR, Cat# 27534.187), all with 5% goat serum, as listed (SI Appendix Table 2). Note that we chose antibodies against Aβ to provide good coverage of the pre-plaque aggregation steps, including Aβ42 monomers/dimers (IBL)(71), Aβ prefibrils (A11)(72) and Aβ protofibrils (OC)(73). Processed tissue was mounted on Superfrost™ glass slides (Thermo Fisher Scientific) from a solution of 50 mM tris(hydroxymethyl)aminomethane (Millipore, Cat# 1.08382.1000) with hydrochloric acid, at pH 7.6, and then left to dry overnight before being coverslipped using entellan (Merck KGaA, Cat# 1.07960.0500). Due to a technical issue, we did not obtain reliable data for the second series of IHC-experiments (second set of experiments) involving reelin and Aβ42 monomers/dimers (IBL).

### Proximity ligation assay (PLA)

We based our PLA protocol on that for Aβ42 and reelin as described above, with the following adaptations according to the manufacturer’s description (Sigma-Aldrich, DUO92101/DUO92103/DOU92012): brain sections were rinsed three times for 10 minutes in PB, then five times for 10 minutes in TBS (50 mM Tris, 150 mM NaCl, pH 8.0) and pre-incubated for 1 hour in 10% normal goat serum in TBS-TX (solution of 50 mM Tris, 0.87% sodium chloride and 0.5% Triton X-100). Then, sections were incubated with primary antibodies (IBL Aβ42 and reelin, see SI Appendix Table 2) in 10% normal goat serum in TBS-TX for 48 hours. After rinsing, we blocked endogenous peroxidase activity by incubating the tissue in hydrogen peroxide solution for 10 min at room temperature, followed by rinsing in “washing solution A”. The tissue was then mounted onto glass slides. Secondary probes attached to oligonucleotides were then added, and, after rinsing, the oligonucleotides of the bound probes were ligated, amplified and visualized by addition of the detection reagent and substrate solution. We then added the nuclear stain solution provided by the kit, followed by dehydrating the sections by immersing them in increasing concentrations of ethanol, ending with a defatting step in xylene for 5 minutes. For each PLA-experiment, we ran sets of technical negative controls. These included omission of either or both primary antibodies, omission of either or both PLA probes, and omission of all the above.

### Imaging and digital processing

We scanned all tissue at 20x using a Zeiss Axioscanner (Z.1) with Zen software (2.6, Blue Ed.) under identical settings (488 channel for EGFP viral tag; 546 channel for Aβ/APP; 635 channel for reelin), taking care to determine the optimal dynamic range for our labeled sections. We identified alEC LII (specifically, DLE and DIE) using established criteria(74). For all sections with infected neurons in dlLEC we used the Zen Circle Tool to place a circle around each EGFP-positive neuron with a diameter of ∼20 µm located in the superficial part of layer II (sometimes referred to as layer IIa). This approach ensures reliable inclusion of Re+ alEC LII principal neurons, while avoiding calbindin neurons that are situated deeper (LIIb). We then read out the mean pixel intensity for the 546 and 635 channels of each EGFP-positive neuron in Zen. To measure non-infected neurons, we took the contralaterally matching level of alEC (same distance from the rhinal sulcus) on each section, and used the same circle tool on Re+ neurons (i.e. the 635 channel). We obtained background levels for each animal independently by using measurements of the cerebellar white matter; these measurements were subtracted from the value of each neuron of each animal. We then exported the data to Python Pandas DataFrames to carry out the statistical analyses. Note that animal 24868 immunolabeled against A11 and reelin was excluded due to a technical error.

### Statistical analysis of differences in protein levels between conditions

We considered the fluorescence level (average pixel intensity) of the protein (reelin vs forms of iAβ), obtained as described in section *Imaging and digital processing*, of all selected neurons within one animal from both hemispheres (conditions: experimental vector-infected and control vector-infected; experimental vector-infected and non-infected; control vector-infected and non-infected). We then subtracted the background level of the protein for each animal. A few negative values occurred, and these were set to zero. Given the substantial biological variability in the physiological levels of reelin and iAβ(38, 39), data from each single animal and both conditions were normalized, by linearly mapping the fluorescence level in the [0,1] interval(75, 76). The normalized data from all animals were then pooled together, while keeping track of the condition. We thus used the normalized data to perform Bayesian estimation of the parameters of a Student-T distribution for each condition, with the degrees of freedom parameter shared between the two conditions(77). Monte Carlo samples from the posterior distribution of the means (*μ*_1_,*μ*_2_) and standard deviations (*σ*_1_,*σ*_2_) for the conditions 1 and 2 were used to estimate the posterior distribution of the effect size(78), defined as:

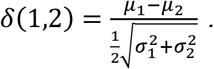

We adopted (−0.2,0.2) as region of practical equivalence for the effect size(79); *δ* ∈(−0.5,−0.2) or (0.2,0.5) was regarded as a small effect size, while *δ* < −0.5 or > 0.5 as big one. For comparison, Bayesian estimation of the effect size was also performed on the normalized data from all animals after randomly assigning the labels of the condition. This analysis was conducted independently on reelin fluorescence data, the three different forms of iAβ, and 1D1.

### Regression analysis of reelin vs iAβ fluorescence levels

For all selected cells fluorescence levels (average pixel intensity), obtained as described in section *Imaging and digital processing*, were normalized at the single protein and animal level, as explained in section *Statistical analysis of differences in protein levels between conditions*. After pooling together the normalized data across animals in each single condition (experimental vector-infected, control vector-infected, non-infected), we performed Bayesian non-parametric regression(80, 81) treating the reelin level as the independent variable “x” and the level of each single *iAβ* form as the dependent variable “y”. The likelihood *p*(*y*|*x*) is normal, with a stationary standard deviation ε, while the mean of the Gaussian is modelled as a function of the independent variable *f(x)*. For the mean function we employed a Gaussian process prior with exponential quadratic kernel, which assumes smoothness but allows for a wide range of functional relationships between x and y(82).

Finally, we sampled from the joint posterior of *f(x)* and ε to build a distribution of the mutual information

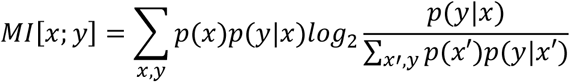

between the dependent variable and the independent one(83). Only for computing mutual information, “x” was discretized over the range covered by the normalized data for that specific condition, and the probability of “x”, *p*(*x*),assumed to be flat.

For comparison, Bayesian estimation of mutual information between reelin and iAβ was also performed on the normalized data for a specific condition from all animals after randomly pairing reelin levels to iAβ levels. Bayesian regression was conducted independently on the three different forms of iAβ and 1D1 from each hemisphere (condition).

### Analysis of EC hippocampal afferents

The fact that infected Re+ alEC LII-neurons express our viral constructs (EGFP) also in their axons allows for clear identification of their terminals in the outer molecular layer of DG (omlDG). Thus, to quantify levels of reelin and Aβ in these terminals from experimental vector-infected vs control vector-infected sides, we used each available section containing omlDG from the dorsal 1/3 with EGFP-positive terminals (SI Appendix Table 3), acquired 16-bit images of each (under identical settings), and exported these to Fiji (ImageJ 1.53i). In Fiji, we programmed a macro to carry out the following automated processing for each image: (1) identify EGFP terminal plexus in omlDG, (2) threshold against background and generate a mask of terminal plexus, (3) apply this mask separately on channels 546 and 635, (4) quantify average pixel intensity within the mask for each channel.

## Results

To test whether *the selective vulnerability of Re+ alEC LII-neurons to accumulate iAβ is linked to their expression of reelin*, we designed a comprehensive set of experiments in which we assessed the effects on levels of iAβ of lowering the expression of reelin in Re+ alEC LII-neurons of AD-rats with the use of a viral vector expressing Re-miRNA (Figure 1).

**Figure 1.**
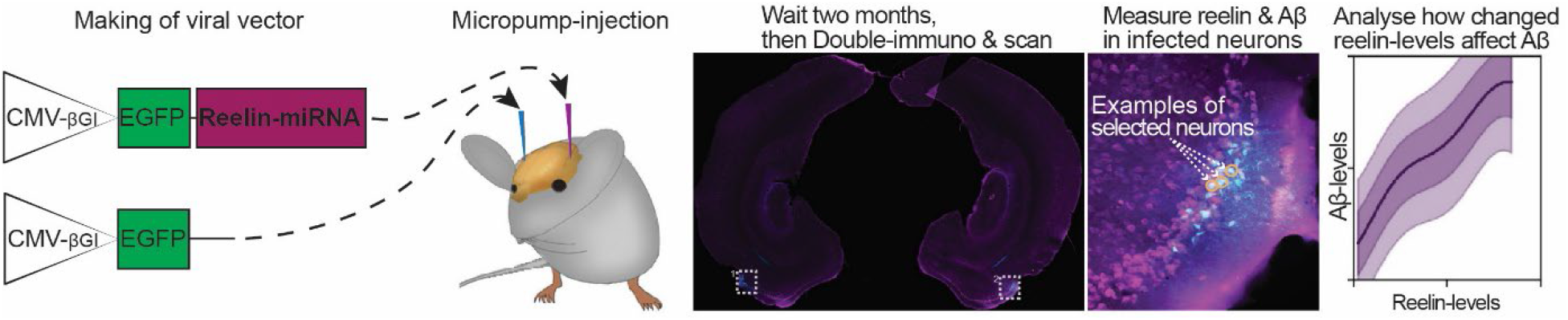
Overview of experimental setup to lower reelin in reelin positive neurons in layer II of the anterolateral entorhinal cortex (alEC) of AD rats. We used an in-house made AAV that expresses miRNA-Re to lower levels of reelin in LII neurons in alEC (dorsolateral LEC). The control virus was similar, though lacked the miRNA-Re. Tissue sections from all animals were stained for the presence of reelin and three different forms of iAβ (Aβ42 with IBL, prefibrils with A11 and protofibrils with OC). Levels of fluorescence were quantified using densitometry on automatically scanned sections and the obtained data were normalized and statistically assessed using Bayesian estimation of the parameters of a Student-T distribution for each condition.

### Reelin directly interacts with Aβ in Re+ alEC LII-neurons

We first immunolabeled sections of three-month-old AD-rats with three different antibodies, selected to capture Aβ across a wide range of configurations prior to formation of mature fibrils that are Aβ plaque-associated. We used a C-terminal specific Aβ42 antibody that detects monomers/dimers (IBLAβ42)(71), a conformation-dependent antibody that detects Aβ prefibrils (A11)(72), and a second conformation-dependent antibody that detects Aβ protofibrils (OC)(73). Notably, the binding of prefibrillar forms of Aβ by A11 is mutually exclusive to the binding of protofibrillar forms of Aβ by OC(73). In alEC, each of these three antibodies react with intracellular material that is selectively present in Re+ LII-neurons, and we observed that each of these forms of Aβ co-localize with reelin to a high degree (SI Appendix, Fig. 1 A-C). This corroborates and extends earlier findings about Re+ alEC LII-neurons where a different set of antibodies against reelin and Aβ were used(39).

To substantiate these confocal results, we used proximity ligation assay (PLA) since this is a highly reliable method to reveal if two proteins are closer than 40 nm(84). We found clear and numerous PLA-signals in superficially located alEC LII-neurons in AD rats at pre-plaque stages, demonstrating a direct interaction between reelin and Aβ42 in these neurons (Fig. 2A). Because neurons under normal conditions make low amounts of Aβ, including Aβ42(85), we decided to test whether a reelin-Aβ42 interaction is detectable also in wild-type rats. Notably, using the same Aβ42 and reelin antibodies as before, on age-matched wild-type Wistar rats, we again find alEC LII neuron-restricted PLA-signals, albeit in very low numbers (Fig. 2B). This latter finding indicates that the tendency of selective accumulation of iAβ42 in Re+ alEC LII-neurons is not an artefact of the transgene-expression of the AD rat model but likely represents a cell-biological feature of Re+ alEC LII-neurons.

**Figure 2.**
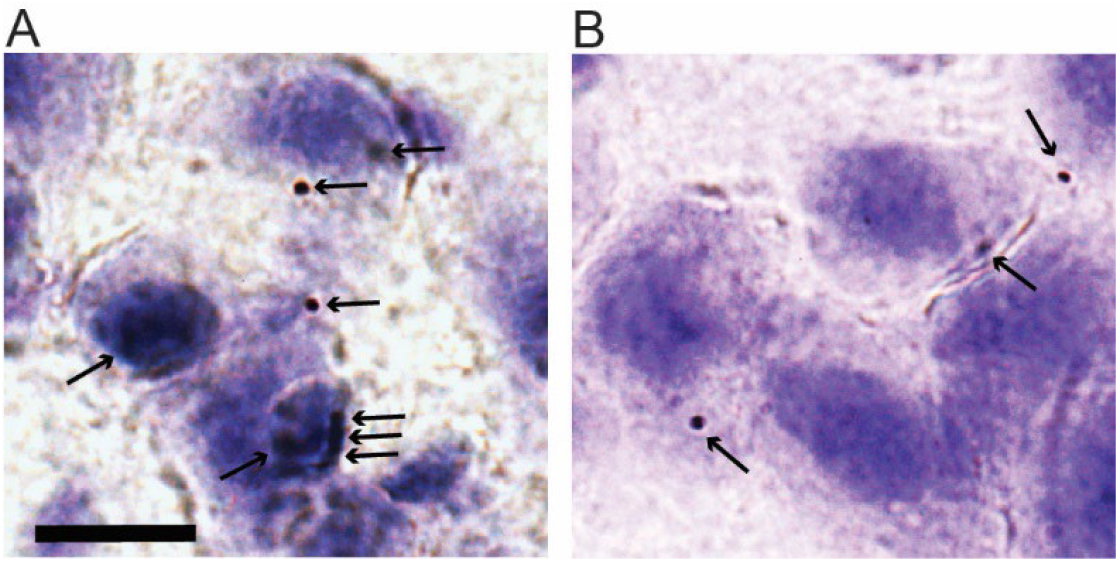
Proximity ligation assay reveals protein-protein interaction between reelin and intracellular Aβ42 in reelin positive neurons in layer II of the entorhinal cortex. **(A)** High powered (100xOil) brightfield micrograph of a representative cluster of alEC layer II-neurons from McGill AD rat reveals multiple dark red spots (arrows), signaling that intracellular Aβ42 and reelin are within ∼40 nanometer of each other. **(B)** We find occasional signals also in layer II-neurons from normal (non-AD) littermate controls, confirming that the dramatically increased incidence seen in AD animals reflects their intracellular Aβ42 accumulation. The counterstain is Cresyl-Violet (Nissl-staining). Scalebar in A, applicable to B as well, equals 10μm.

### Lowering Reelin in Re+ alEC LII-Neurons Concomitantly Reduces iAβ

We next set out to test whether a functional interaction exists between the two proteins. To selectively lower reelin in Re+ alEC LII-neurons, we constructed an *experimental vector* carrying miRNA against reelin (CMB-βGI-EGFP-Reelin-miRNA) plus a *control vector* (CMB-βGI-EGFP). We successfully placed selective bilateral injections, the experimental vector on one side and the control vector on the contralateral side, into LII of alEC in five AD rats (Fig. 1; SI Appendix, Fig 2). To target the earliest possible AD-related stage of the model, after EC has fully developed, we injected animals at ages a few days past 1 month(86).

Confocal images of a randomly selected sets of immunolabeled Re+ alEC LII-neurons expressing the experimental vector were compared with images of Re+ alLII-neurons at the corresponding dorsoventral position of the contralateral alEC expressing the control vector. The experimental vector efficiently lowered reelin-levels relative to the control vector. Furthermore, the immunostaining against iAβ revealed a concomitant reduction for each of the three aggregation states tested, i.e., Aβ42 monomers/dimers, Aβ prefibrils and Aβ protofibrils (Fig. 3 A-C).

**Figure 3.**
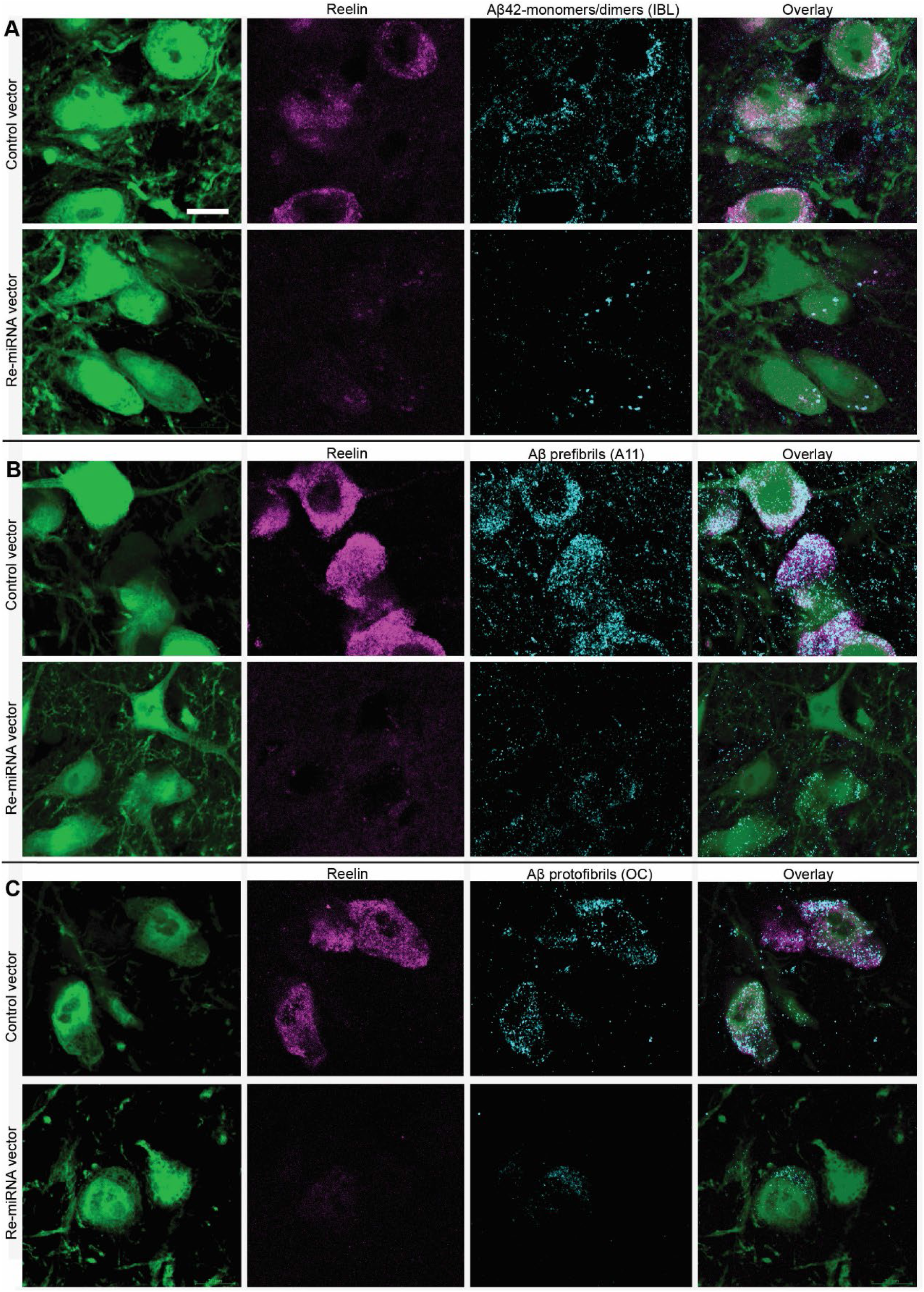
Lowering levels of reelin in Re+ alEC LII-neurons results in a concomitant reduction of levels of iAβ across three aggregation states. **(A-C)**. Each panel shows confocal optical sections of 0.7μm thickness to illustrate the effect of the experimental vector (miRNA-Re) relative to control for reelin and **(A)** Aβ42 monomers/dimers, **(B)** Aβ prefibrils and **(C)** Aβ protofibrils. The micrographs were acquired using identical settings. Scalebar in (A) is for all micrographs and equals 10μm.

To quantify the efficiency of the lowering of reelin and the concomitant reduction of iAβ in Re+ alEC LII-neurons, we used fluorescence densitometry. The average fluorescence levels for reelin and each of the three forms of iAβ (i.e., three different IHC-procedures) for individual neurons infected with the *experimental vector* or the *control vector* was quantified. This quantification showed that the level of reelin was significantly reduced by an amount ranging in effect size from 1.6-2.1 standard deviations for neurons infected by the experimental vector relative to neurons infected by the control vector. This led to a concomitant reduction of iAβ that for Aβ42 monomers/dimers (IBL) amounted to an effect size of 0.93 standard deviations (experimental vector n = 820, control vector n = 585), for prefibrils (A11) the effect size was 0.6 standard deviations (experimental vector n = 685, control vector n = 579), and for protofibrils (OC) the effect size was 0.45 standard deviations (experimental vector n = 365, control vector n = 283; Fig. 4).

**Figure 4.**
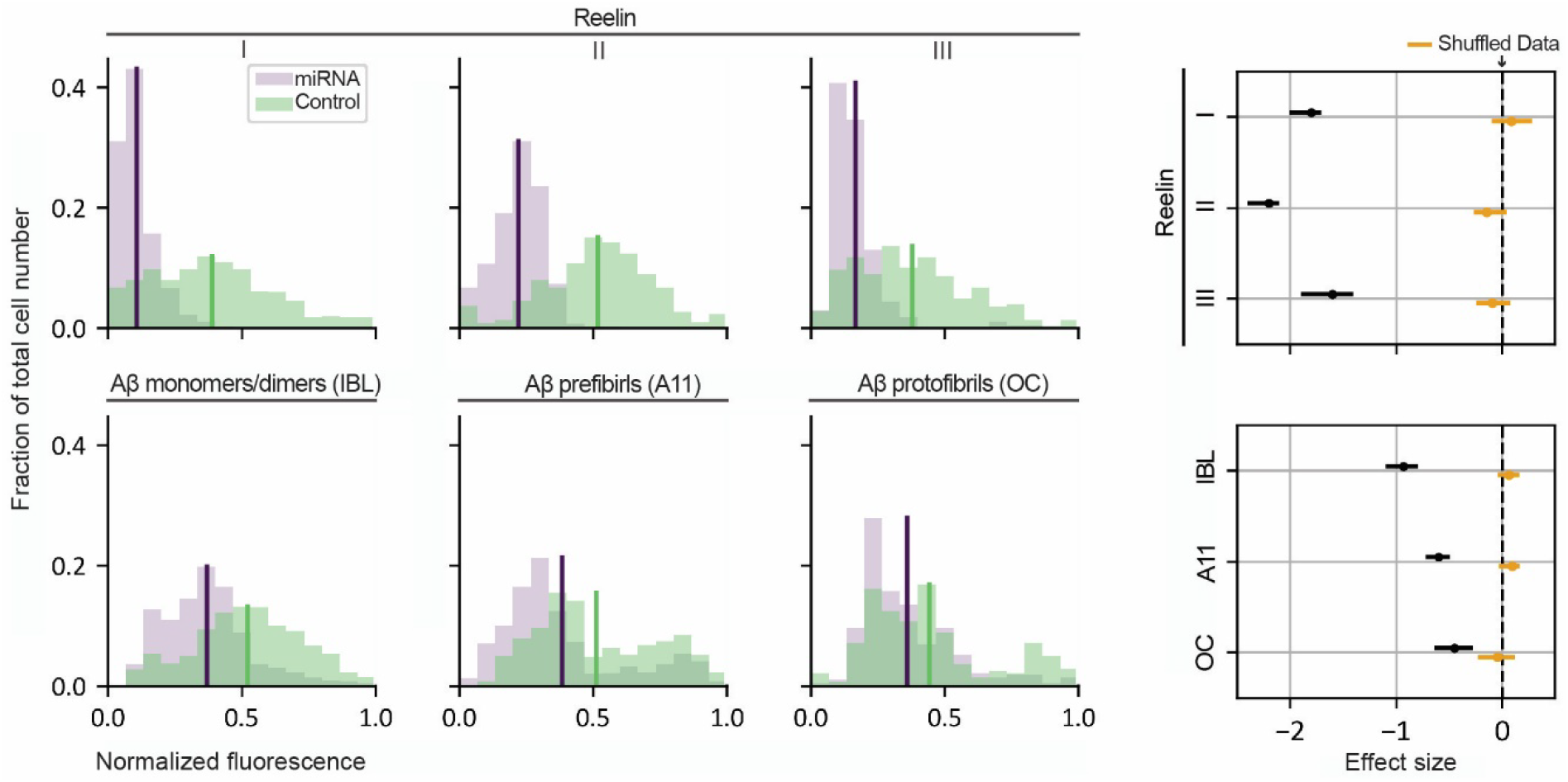
Quantification of the lowering of reelin and the concomitant reduction of iAβ across three aggregation states in Re+ alEC LII-neurons. **(A)** Upper row: plots showing levels of reelin as measured by average immunofluorescence per neuron (binned) following injections of the experimental vector (miRNA-Re) to lower reelin-levels, relative to a control vector. Lower row: plots showing the resulting effects from lowering reelin levels on levels of three different forms of Aβ, including Aβ monomers/dimers (IBL), Aβ prefibrils (A11) and Aβ protofibrils (OC). **(B)** The associated right-side diagrams shows the effect of the manipulation for each plot (effect sizes expressed as standard deviation). Posterior distribution of the mean of the effect size from the shuffled data are colored in yellow. Solid lines span the 95% credible interval, while the dots represent the mean.

We then inferred a predictive distribution of the levels of Aβ given the reelin-level via Bayesian non-parametric regression, using the average fluorescence level measured for each Re+ alEC LII-neuron for each of the three IHC-procedures. This showed that, regardless of whether reelin was lowered or not, the levels of reelin are predictive of the levels of each form of Aβ (monomers/dimers, prefibrils and protofibrils). Bayesian estimation of mutual information indeed reveals a clear dependency between reelin and each form of Aβ, with no overlap between the 95% credible interval for the measurements and the shuffle distribution (Fig. 5).

**Figure 5.**
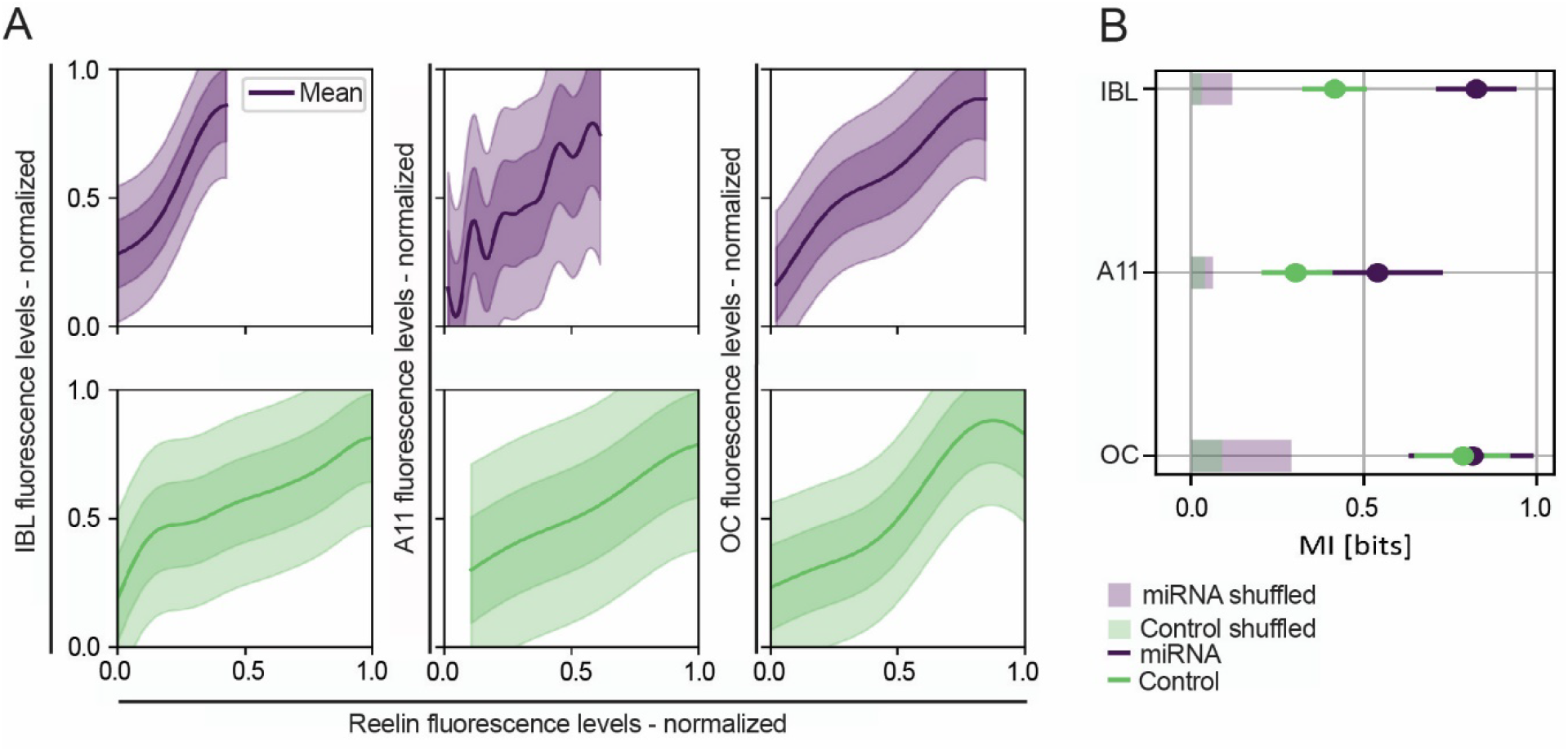
*Mutual information between* Aβ monomers/dimers (IBL), Aβ prefibrils (A11), Aβ protofibrils (OC) and reelin. (**A)** Posterior predictive distribution p(y|x) of iAβ levels (y-axis) vs reelin level (x-axis) (see Methods section, *Regression analysis of reelin vs iAβ fluorescence levels*) for the neurons infected by the experimental vector (miRNA-Re; top, purple) and for the neurons infected by the control vector (bottom, green). Each column corresponds to fluorescence data collected in all the experiments in which a specific form of Aβ was immunolabelled (IBL, A11, OC from left to right). Solid lines mark the mean of the distribution, darker regions the 68%ile of the credible interval and lighter regions the 95%ile of the credible interval. **(B)** Posterior distribution of mutual information MI between levels of each form of Aβ, and levels of reelin (see Methods section, *Regression analysis of reelin vs iAβ fluorescence levels*) for the neurons infected by the experimental vector (miRNA-Re, purple) and for the neurons infected by the control vector (green). Each row corresponds to fluorescence data collected in all the experiments in which a specific form of iAβ was immunolabelled (IBL, A11, OC from top to bottom). Solid lines span the 95% credible interval, while the dots represent the mean. 95% credible interval for the shuffled data are displayed as thick semi-transparent lines.

We previously reported that levels of Aβ vary in a non-systematic way between the left and right EC of the rat model we use(68). We therefore also tested the effects of the experimental vs control vector against neurons in non-injected hemispheres, by placing unilateral injections of either the experimental (9 animals) or the control vector (10 animals) in LII of alEC. Confocal images again showed that the experimental vector efficiently lowered reelin-levels relative to the control vector. The amount was quantified by fluorescence levels for reelin vs prefibrils (A11) and protofibrils (OC) in Re+ alEC LII-neurons infected with the experimental vector, compared with uninfected Re+ neurons from the corresponding dorsoventral position of the contralateral alEC. As the uninfected neurons do not express GFP, we selected these by their expression of reelin. We used the same approach for animals injected with the control vector. For this second set of experiments, the experimental vector reduced reelin by 2.1 standard deviations. This led to a concomitant reduction of levels of Aβ prefibrils (A11) with an effect size of 0.6 standard deviations (experimental vector n = 1601, non-infected n = 2059), and for levels of protofibrils (OC) the reduction had an effect size of 0.32 standard deviations (experimental vector n = 1980, non-infected n = 1892), thus substantiating the findings from the first set of experiments. The control vector led to a reduction of reelin amounting to 0.45 standard deviations relative to uninfected neurons. This had no impact on the levels of Aβ (prefibrils, control vector n = 2244, non-infected n = 2219; protofibrils, control vector n = 1178, non-infected n = 1180; SI Appendix, Fig. 3).

The inferred predictive distribution of the Aβ levels (A11 and OC) given the reelin levels substantiated our findings from the first set of experiments, thus showing that the levels of reelin are predictive of the levels of Aβ prefibrils (A11) and Aβ protofibrils (OC). Bayesian estimation of mutual information reveals a clear dependency between reelin and both forms of Aβ, with no overlap between the 95% credible interval for the measurements and the shuffle distribution (SI Appendix Fig. 4).

### The Reduction of iAβ by Lowering of Reelin in Re+ alEC LII-Neurons is Independent of APP-levels

The transgene of the rat model we used drives expression of mutated hAPP, and these mutations shift the processing of the hAPP to cause pathological amounts of Aβ(45). Therefore, any manipulation that changed Aβ-levels could stem from changes in hAPP-levels. To assess whether the effect of lowering reelin upon iAβ resulted from changes in hAPP-levels, we used a well-validated hAPP antibody (1D1)(87) on tissue from both the first and the second set of experiments. To check whether measurements might be influenced by the use of fluorophores or chromogenic labeling, we used immunoenzyme labeling with 3,3’-Diaminobenzidine (DAB) for the material from the first set of experiments, and fluorophores (as used for the above IHC) for the second set of experiments. The results from the first set of experiments using DAB showed that neurons infected with the experimental vector retained the same levels of hAPP as neurons infected with the control vector. Data from the second set of experiments substantiate this, as neurons infected with the experimental vector showed only a minor reduction of hAPP-levels relative to neurons infected with the control vector, amounting to 0.15 standard deviations (Numbers of neurons for first set of experiments: experimental vector n = 237, control vector n = 156. Numbers of neurons for second set of experiments: experimental vector n = 1809, non-infected n = 1798; control vector n = 2196, non-infected n = 1682; Fig 6). Bayesian non-parametric regression shows that, contrary to the case for reelin-levels upon Aβ-levels, reelin-levels are not predictive for levels of hAPP. Results from Bayesian estimation of mutual information furthermore indicates a lack of dependency between reelin and hAPP (SI Appendix Fig. 5).

**Figure 6.**
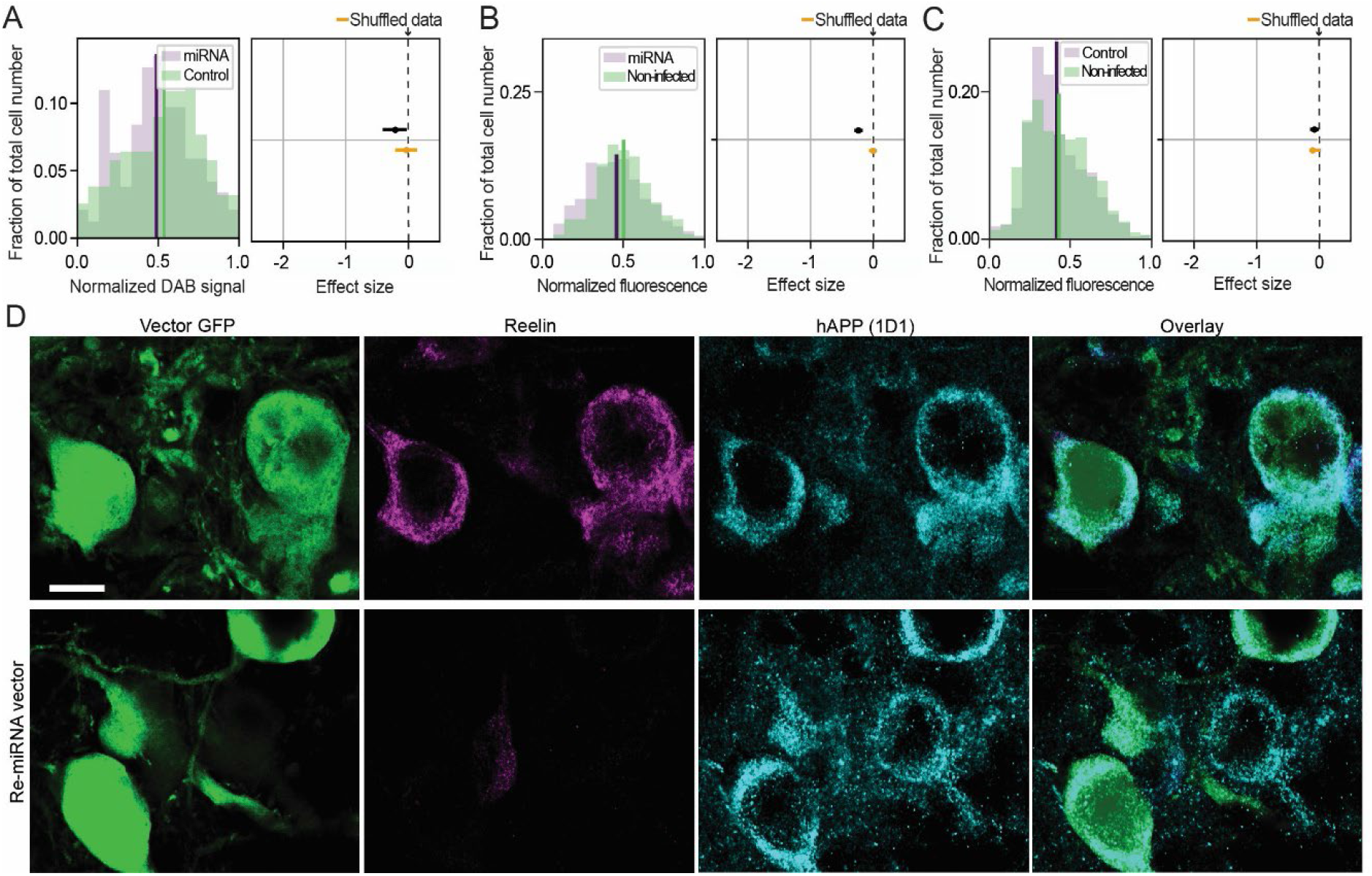
Reduced accumulation of iAβ, induced by lowering reelin in Re+alEC LII-neurons, occurs without substantial associated changes in human APP-levels. **(A)** Double-injected animals: Left-side plot shows levels of human amyloid precursor protein (hAPP, antibody 1D1), measured using DAB, in neurons infected with the experimental vector (miRNA-Re) to lower reelin-levels, compared with contralateral neurons infected with a control vector. Right-side plot shows that the experimental vector does not have any effect on levels of hAPP (effect size expressed as standard deviations (SD)). **(B)** Single injected animals: Left-side plot shows levels of hAPP (1D1), measured using fluorescence, in neurons infected with the experimental vector (miRNA-Re) to lower reelin-levels (10 animals), compared with non-infected neurons. Right-side plot indicates that the experimental vector has a minor effect on levels of hAPP (effect size = −0.15 SD). **(C)** Same as for (B) but for neurons infected with a control vector (9 animals) and compared to non-infected neurons. Right-side plot shows that the control vector does not have any effect on levels of hAPP. **(D)** Double-immunofluorescence of Re+ alEC LII-neurons in single injected animals. Top row: Confocal optical sections of 0.7μm thickness illustrate that upon infection with the control vector, reelin is readily apparent along with human hAPP (1D1). Bottom row: infection with the experimental vector (miRNA-Re), which effectively reduces reelin, does not lead to a visible change in levels of hAPP. The micrographs were acquired using identical settings. Scalebar in top left micrograph represents all and equals 10μm

### Lowering Reelin in Re+ alEC LII-Neurons reduces levels of reelin and oligomeric Aβ in hippocampal terminals

Having found that lowering Reelin in Re+ alEC LII-neurons concomitantly reduces the levels of iAβ in their somata, we next examined whether this affects reelin and/or Aβ in the area covered by the hippocampal terminal plexus of these neurons in the dentate gyrus and CA3/2. To this end, we performed automated densitometric quantification of the fluorescence levels of reelin plus Aβ prefibrils (A11) and Aβ protofibrils (OC) in the area covered by the labelled hippocampal plexuses. We used every section with a GFP-labeled plexus in the dentate gyrus/CA3 from the animals of the second dataset (single injected animals; number of sections from each animal is listed in SI Appendix Table 3). Lowering Reelin in Re+ alEC LII-neurons reduced levels of reelin in the hippocampal terminals of these neurons by 33%, and this led to a concomitant 37% reduction of prefibrilar Aβ (SI Appendix Fig.6). Meanwhile, we found a non-significant trend towards reduction for protofibrillar Aβ (data not shown).

## Discussion

In a previous paper, we showed that Re+ neurons, a substantial and specific subset of principal neurons in alEC LII, are prone to accumulate iAβ(39). Our current data show that long before Aβ-plaques start to form, reelin and iAβ42 can structurally associate in these neurons. We further show that selectively lowering the amount of reelin in the Re+ alEC LII neurons reduces their propensity to accumulate iAβ. This reduction encompasses three levels of Aβ-aggregation, monomers/dimers, prefibrils, and protofibrils, and occurs without any substantial associated changes in hAPP-levels.

Aβ-plaques no doubt play a major role in the context of full-blown AD, but the situation is different in the prodromal phase of the disease. For example, studies on living human subjects show that cerebrospinal fluid-levels of Aβ, as measured with flow cytometry using microsphere-based Luminex xMAPTM technology, change before formation of Aβ-plaques, where the latter were revealed by positron emission tomography(10, 27). Biochemical studies align with this by showing that in the human brain, levels of non-fibrillated Aβ predict cognitive decline and neurodegeneration better than the amount of Aβ-plaques(32-37). This has been corroborated by experimental results obtained in animal models(38, 40, 41) and neuron cultures(38, 42). These findings are further substantiated by human post-mortem studies using sensitive immunohistochemical methods, showing that Aβ starts to accumulate intracellularly prior to formation of Aβ-plaques(28-31), and that such iAβ-accumulation is striking in EC LII-neurons(31, 39, 88-90). This body of evidence provides a strong argument that non-fibrillated forms of Aβ are highly relevant in the onset of AD(91).

The reelin-Aβ interaction we report here corroborates earlier reports that the two molecules may directly interact in the brain as well as in cultured cells(59, 60). These interactions probably depend on molecular features of the two molecules. Reelin is a self-associating and cysteine-rich protein that contains multiple β-strands(92, 93). Alongside the potential for β-strands to aggregate, it is worth noting that Aβ-peptides exhibit pathological binding specifically to cysteine-rich domains(94). Another relevant factor is that the reelin C-terminus, which promotes binding and helps to induce downstream signaling events, has a strong positive charge(95), and Aβ42-peptides have a negative charge(96). Recent findings show that neuron-specific increases of proteins that exceed their solubility-levels will drive these proteins towards aggregation(44). Such an exceeding of solubility-levels is likely at play also regarding findings that Aβ causes a dose-dependent increase of non-signaling competent reelin. The latter observations were made in cell-cultures and substantiated by findings in the human Aβ-affected neocortex(58, 59), where reelin and oligomeric Aβ were shown to co-immunoprecipitate(60). All of these factors can be joined in the context of Re+ alEC LII-neurons and so help explain their selective accumulation of iAβ.

The interaction between reelin and Aβ prevents reelin from triggering processing-and internalization of ApoER2(54, 59, 60). This ApoER2 processing normally creates an N-as well as a C-terminal ApoER2-fragment, both detectable in cerebrospinal fluid. In line with the Aβ-induced impairment of reelin signaling found in cultured cells, the amounts of these ApoER2-fragments drop by about 50% in CSF of AD-subjects, even though levels of *ApoER2 messenger RNA* as well as full-length ApoER2 remain unaltered(60, 97). This provides convincing corroborative evidence that an interaction between reelin and Aβ occurs in the brain of AD-subjects, and that this impairs the reelin signaling-cascade. Because this signaling cascade works to enhance glutamatergic transmission in the adult brain and promotes LTP(51, 52), loss of reelin in adulthood lead to deficits in memory functions in mice and monkeys. Such deficits were indeed shown by recent experiments on reelin conditional knockout mice(55), and this is further corroborated by findings that an age-related decline in reelin-levels in Re+ alEC LII-neurons in monkeys associates with memory impairments(98).

Besides its role in memory, reelin-signaling through ApoER2 initiates a second signaling-cascade mediated by tau-phosphorylating kinases. When initiated, this cascade potently inhibits the activity of one of the main tau-phosphorylating kinases, namely GSK3β(61-63). Metabolically active reelin thus regulates the activity of GSK3β such that low levels of reelin result in increased levels of p-tau, as demonstrated in congenitally reelin-deficient mice (reeler-mice) that show a strong increase of p-tau(61). A converse situation is apparent when mating transgenic mice overexpressing reelin, with transgenic mice expressing mutated human tau that normally develop ample p-tau by 6 months. The resulting offspring exhibit a dramatic reduction of p-tau(76). Furthermore, the application of reelin-Aβ aggregates onto mouse primary neurons result in a two-fold increase in levels of p-tau in the cell-extracts relative to that seen following the addition of pure reelin (59). These converging results convincingly show that increased reelin-signaling serves to downregulate the phosphorylation-state of tau. Vice versa, one might thus predict that lowered levels of active reelin, resulting from the molecular interactions between reelin and iAβ42, will result in an increased phosphorylation of tau. This is of interest in view of the strong evidence that the predominant cortical onset of NFTs occurs in alEC LII (11-15). Moreover, in the rat model, levels of iAβ and reelin in EC LII show a clear gradient, with both being high in neurons situated anterolaterally, but gradually lower in neurons situated successively more medially(39). Future work should address whether these graded levels of iAβ and reelin relate to the gradient established for NFTs, which, following their onset in alEC LII, progressively invade more medially situated LII-neurons(11, 13, 99).

The above findings, alongside our present findings of a pre-plaque interaction between reelin and Aβ in Re+ alEC LII-neurons, leads us to propose the following model as to why these neurons form NFTs in prodromal AD (Fig. 7). It has been argued that alEC LII-neurons have a very high metabolic rate, and, as the autophagic machinery becomes less efficient owing to ageing, iAβ will tend to accumulate at a disproportional rate in these neurons(43). When this happens, iAβ begins to pathologically interact with reelin, causing further accumulation of iAβ, and further increasing the workload upon, and failure of, the autophagic machinery. The interaction of reelin with iAβ will then increasingly lead to signaling-deficient reelin supplanting signaling-competent reelin. This causes a failure of reelin to bind ApoER2 and a consequent downregulation of the signaling cascades normally controlled by reelin, leading to impairments in entorhinal-hippocampal LTP, and, crucially, a failure to downregulate the activity of GSK3β. The subsequent increased activity of GSK3β leads to increased formation of p-tau that finally results in NFTs. The finding that expressing the ε4 variant of apolipoprotein, which constitutes the highest risk-factor for AD aside from ageing, impairs recycling of ApoER2 due to vesicular trapping in the endocytic transport machinery(100), ties in well with our proposed model. This is because one can logically predict that the impaired signaling via the ApoeR2, due to iAβ-induced loss of reelin-signaling, will have a greater impact in subjects in which the ApoER2 is already compromised by expressing the ε4 apolipoprotein variant. This model provides a testable prediction: any compound preventing reelin from associating with iAβ in Re+ alEC LII-neurons, which does not in itself trigger production of more Aβ, will prevent or at least substantially reduce the formation of p-tau and NFTs.

**Figure 7.**
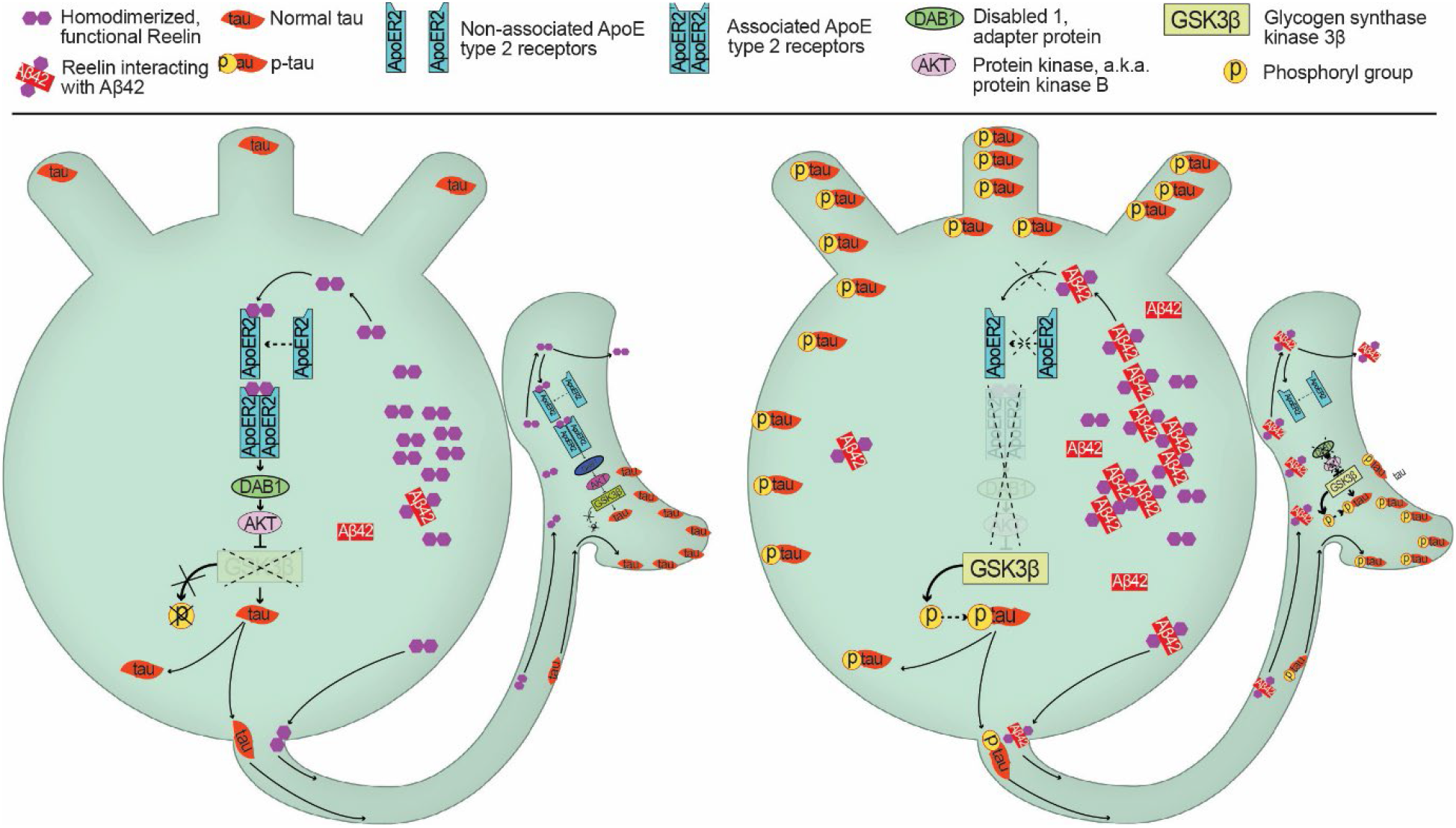
Schematic representation of a mechanistic model for how reelin-iAβ42 interactions lead to a selective vulnerability in formation of p-tau in Re+ alEC LII-neurons. Left-side cartoon: in a healthy Re+ alEC II-neuron there is little accumulation of iAβ42. Intact reelin-signaling leads to a low activity-level of GSK3β and helps ensure the appropriate phosphorylation-state of tau. This is part of the requirement for tau to remain localized to the proper compartments, with the highest levels being in the axon. Right-side cartoon: as the autophagic machinery becomes less efficient owing to ageing, iAβ42 will tend to accumulate at a disproportional rate in Re+ alEC LII-neurons, due to the high metabolic rate of these neurons. When this happens, iAβ begins pathologically interacting with reelin, further increasing the workload upon, and failure of, the autophagic machinery. The interaction of reelin with iAβ will cause reelin-iAβ42 complexes to form and increasingly lead to signaling-deficient reelin supplanting signaling-competent reelin. The reelin-iAβ42 complexes are unable to bind the ApoER2-receptor to initiate the downstream signaling cascade. This removes the inhibitory signaling upon GSK3β, causing the latter to drive hyperphosphorylation of tau (p-tau), which is misallocated away from the axon and into somatodendritic compartments where NFTs form.

## Appendix: Supplementary Information

**Supplementary figure 1.**
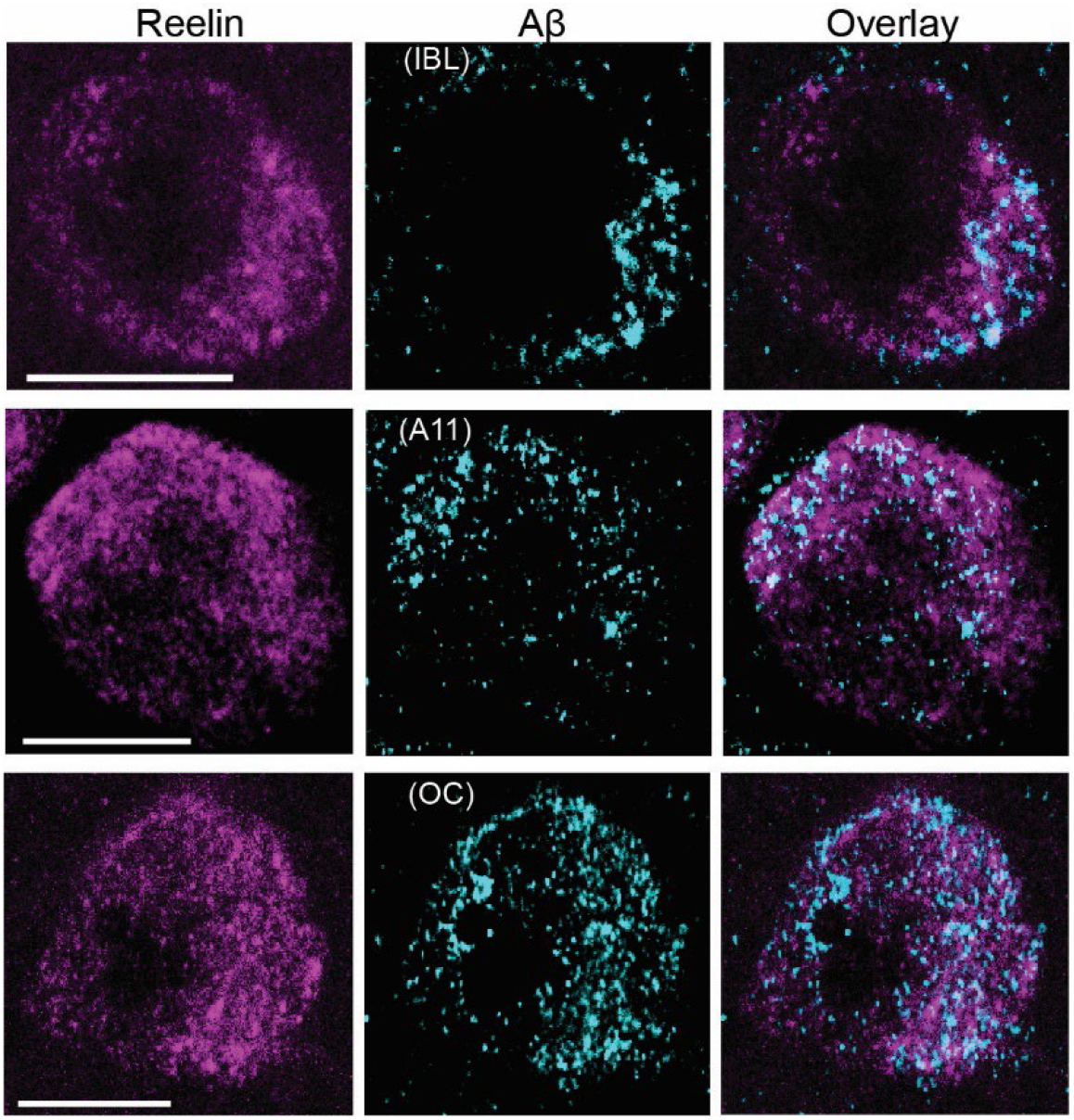
Subcellular co-localization between reelin and different forms iAβ in cytoplasmic granules of reelin positive anterolateral entorhinal cortex layer II-neurons in young adult AD rats. Confocal micrographs of 0.7μm optical sections of double immunohistochemical stains with reelin (left-side micrographs) vs three forms of Aβ (middle micrographs), including iAβ42 monomers/dimers (IBL; top), iAβ prefibrils (A11; middle), and iAβ protofibrils (OC). The right-side micrographs show the overlays. Scalebars = 10μm

**Supplementary figure 2.**
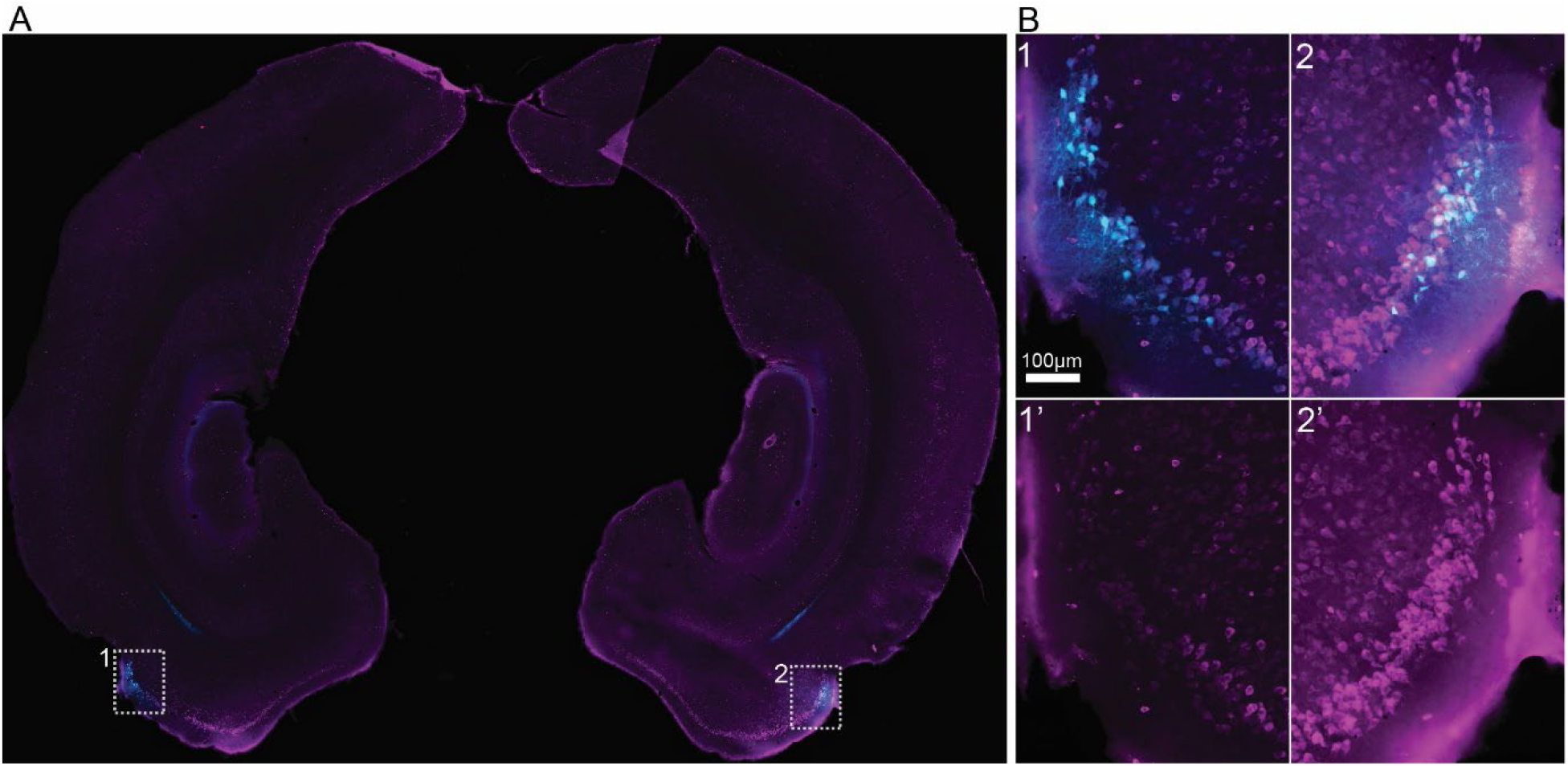
Example of double-injected brain. **(A)** The right anterolateral entorhinal cortex was injected with the control vector, while the left anterolateral entorhinal cortex was injected with the experimental vector (miRNA-Re). Both injection sites and the associated transfection of neurons are labeled due to EGFP-expression (pseudo-colored cyan). **(B)** Higher power image of the insets indicated in A, showing the restriction of the injection (cyan) to the outer part of EC LII (LIIa), and the effect of the experimental vector on the expression of reelin (magenta fluorescence). All imaging settings are identical. Scalebar in (B) = 100 µm.

**Supplementary figure 3.**
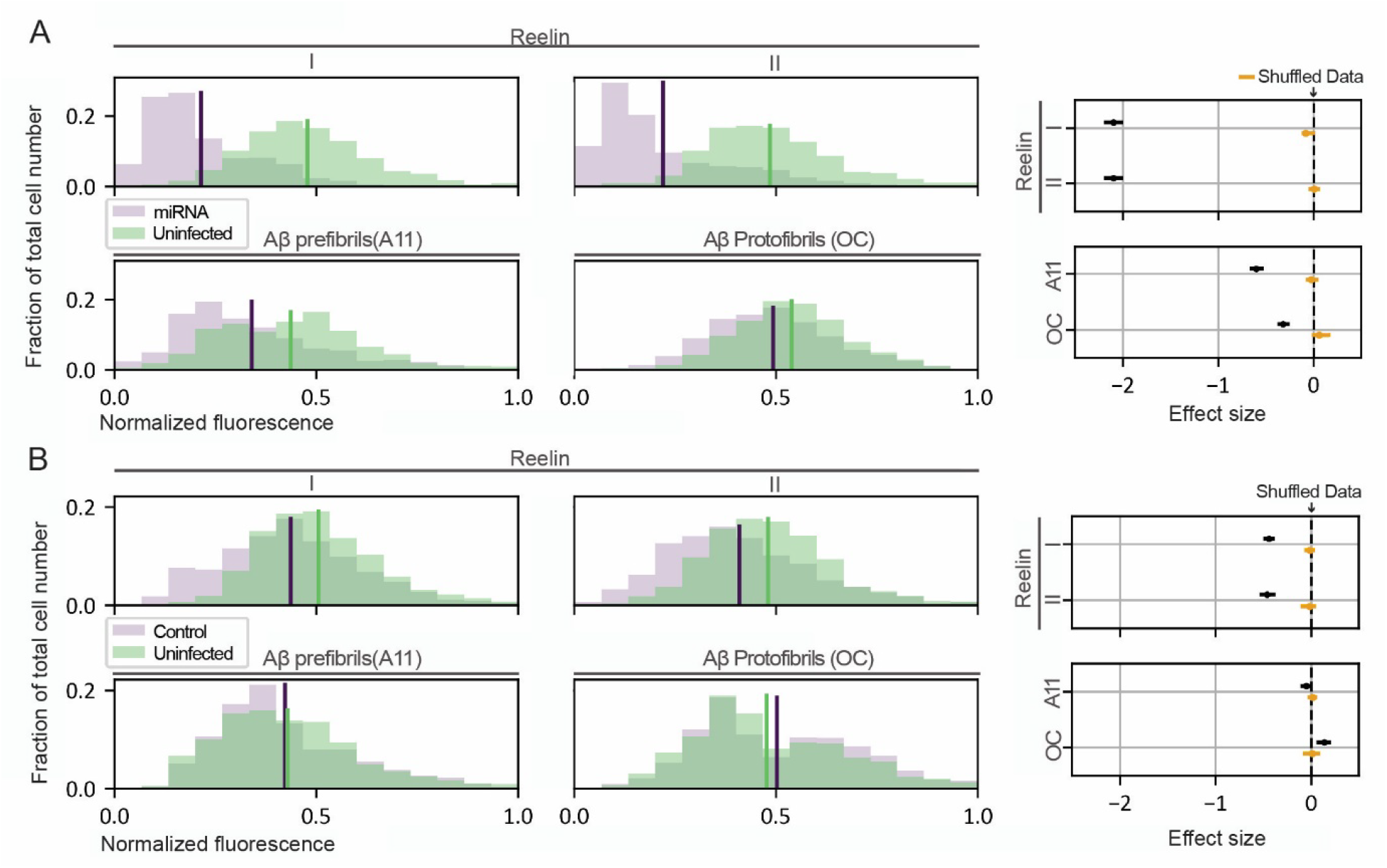
Quantification of the lowering of reelin and the concomitant reduction of iAβ in Re+ anterolateral entorhinal LII-neurons (alEC LII) for the second set of experiments. **(A)** Upper row: plots showing lowered levels of reelin as measured by average immunofluorescence per neuron (binned) following injections of the experimental vector (miRNA-Re) to lower reelin-levels, relative to uninfected Re+ neurons in the contralateral alEC LII; right hand panel shows the efficacy of the manipulation expressed as effect sizes of the standard deviation. Lower row: lowering reelin levels leads to a concomitant lowering of iAβ prefibrils (A11) and protofibrils (OC); right hand panel shows the efficacy of the manipulation expressed as effect sizes of the standard deviation. Posterior distribution of the mean of the effect size from the shuffled data are colored in yellow. Solid lines span the 95% credible interval, while the dots represent the mean. **(B)** Same as for (A) but following injections of a control viral vector, relative to uninfected Re+ neurons in the contralateral alEC LII. The control vector causes a modest reduction of reelin-levels relative to the experimental vector, but this has no effect on iAβ.

**Supplementary figure 4.**
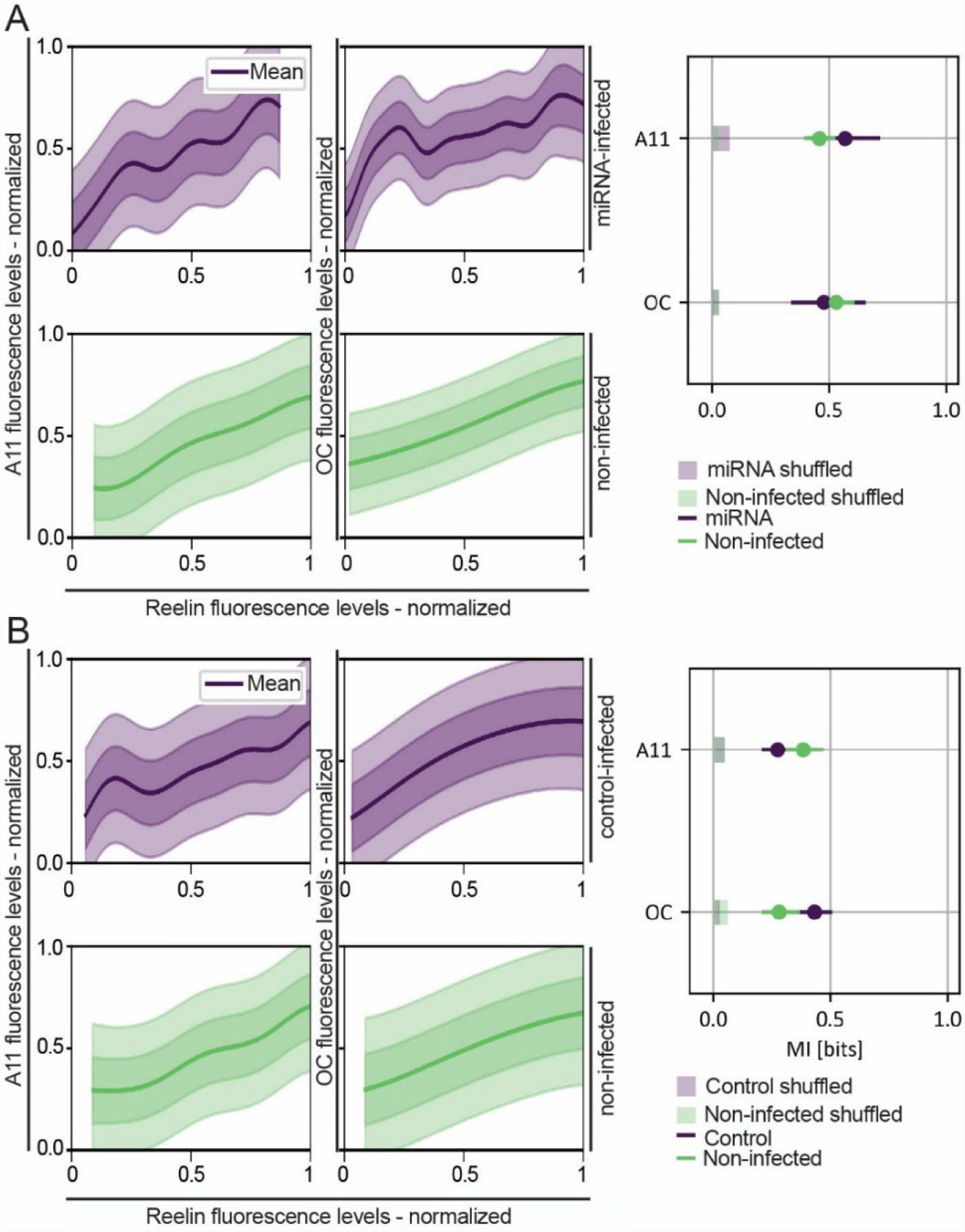
*Mutual information between* Aβ prefibrils (A11), Aβ protofibrils (OC) and reelin. (**A)** Left-side plots: posterior predictive distribution p(y|x) of iAβ levels (y-axis) vs reelin level (x-axis) (see Methods section, *Regression analysis of reelin vs iAβ fluorescence levels*) for the neurons infected by the experimental vector (miRNA-Re; top, purple) and for the non-infected neurons (bottom, green). Each column corresponds to fluorescence data collected in all the experiments in which a specific form of Aβ was immunolabelled (A11, OC from left to right). Solid lines mark the mean of the distribution, darker regions the 68%ile of the credible interval and lighter regions the 95%ile of the credible interval. Right-side schematic: Posterior distribution of mutual information MI between levels of each of the two forms of Aβ, and levels of reelin (see Methods section, *Regression analysis of reelin vs iAβ fluorescence levels*) for the neurons infected by the experimental vector (miRNA-Re, purple) and for non-infected neurons (green). Each row corresponds to fluorescence data collected in both experiments in which the specific form of iAβ was immunolabelled (A11, OC from top to bottom). Solid lines span the 95% credible interval, while the dots represent the mean. 95% credible interval for the shuffled data are displayed as thick semi-transparent lines. **(B)** Left-side plots: posterior predictive distribution p(y|x) of iAβ levels (y-axis) vs reelin level (x-axis) for the neurons infected by the control viral vector (top, purple) and for the non-infected neurons (bottom, green). Each column corresponds to fluorescence data collected in all the experiments in which a specific form of Aβ was immunolabelled (A11, OC from left to right). Solid lines mark the mean of the distribution, darker regions the 68%ile of the credible interval and lighter regions the 95%ile of the credible interval. Right-side schematic: Posterior distribution of mutual information MI between levels of each of the two forms of Aβ, and levels of reelin for the neurons infected by the control viral vector (purple) and for non-infected neurons (green). Each row corresponds to fluorescence data collected in both experiments in which the specific form of iAβ was immunolabelled (A11, OC from top to bottom). Solid lines span the 95% credible interval, while the dots represent the mean. 95% credible interval for the shuffled data are displayed as thick semi-transparent lines.

**Supplementary figure 5.**
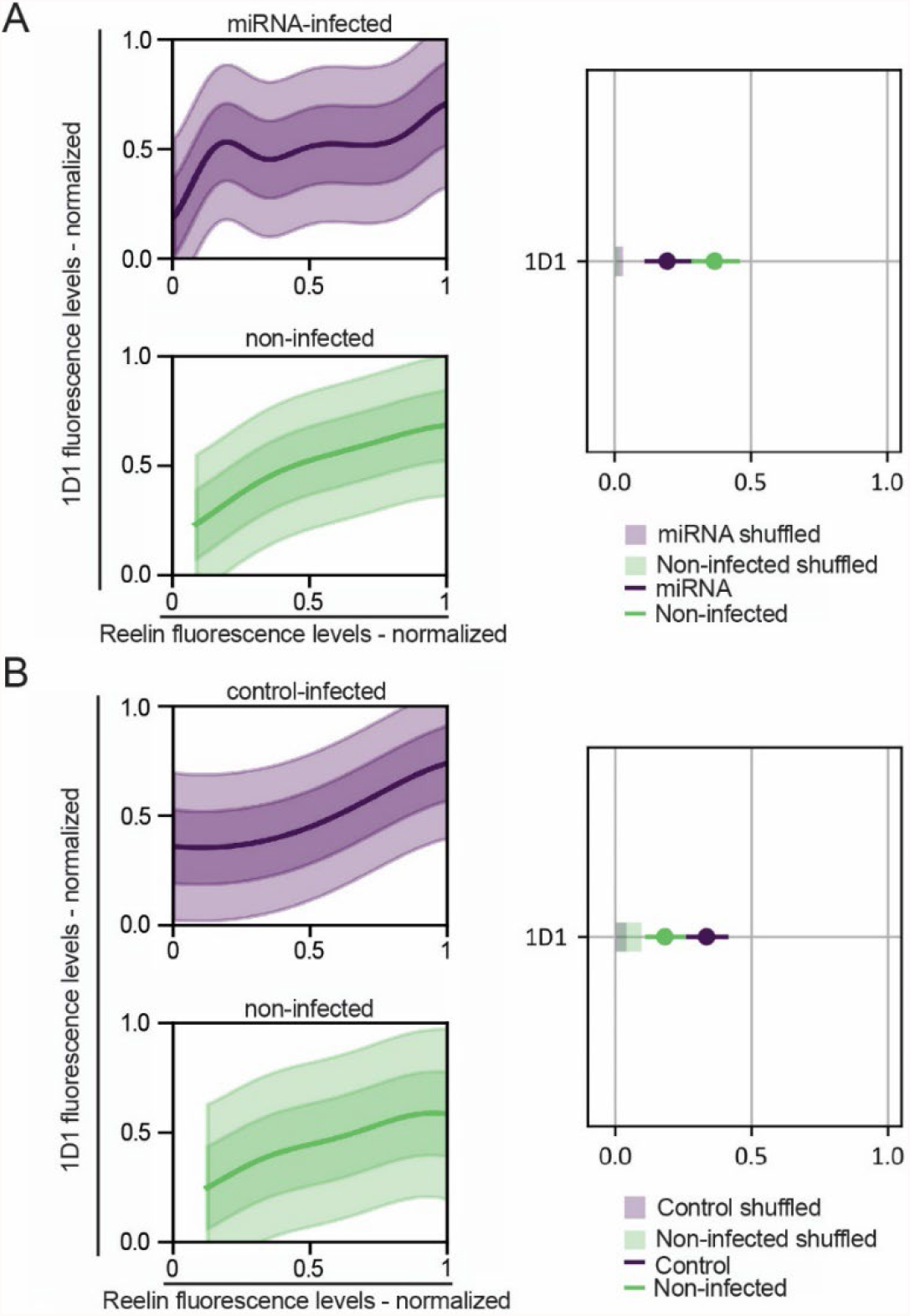
*Mutual information between* levels of human APP (hAPP; 1D1) and reelin, measured from fluorescence-levels. (**A)** Left-side plots: posterior predictive distribution p(y|x) of hAPP (1D1) levels (y-axis) vs reelin level (x-axis) (see Methods section, *Regression analysis of reelin vs iAβ fluorescence levels*) for the neurons infected by the experimental vector (miRNA-Re; top, purple) and for the non-infected neurons (bottom, green). Both columns correspond to fluorescence data collected for the experiments in which hAPP was immunolabelled (1D1). Solid lines mark the mean of the distribution, darker regions the 68%ile of the credible interval and lighter regions the 95%ile of the credible interval. Right-side schematic: Posterior distribution of mutual information MI between levels of hAPP and levels of reelin (see Methods section, *Regression analysis of reelin vs iAβ fluorescence levels*) for the neurons infected by the experimental vector (miRNA-Re, purple) and for non-infected neurons (green). Solid lines span the 95% credible interval, while the dots represent the mean. 95% credible interval for the shuffled data are displayed as thick semi-transparent lines. **(B)** Left-side plots: posterior predictive distribution p(y|x) of hAPP-levels (1D1; y-axis) vs reelin level (x-axis) for the neurons infected by the control vector (top, purple) and for the non-infected neurons (bottom, green). Solid lines mark the mean of the distribution, darker regions the 68%ile of the credible interval and lighter regions the 95%ile of the credible interval. Right-side schematic: Posterior distribution of mutual information MI between levels of hAPP and levels of reelin for the neurons infected by the control vector (purple) and for non-infected neurons (green). Solid lines span the 95% credible interval, while the dots represent the mean. 95% credible interval for the shuffled data are displayed as thick semi-transparent lines.

**Supplementary figure 6.**
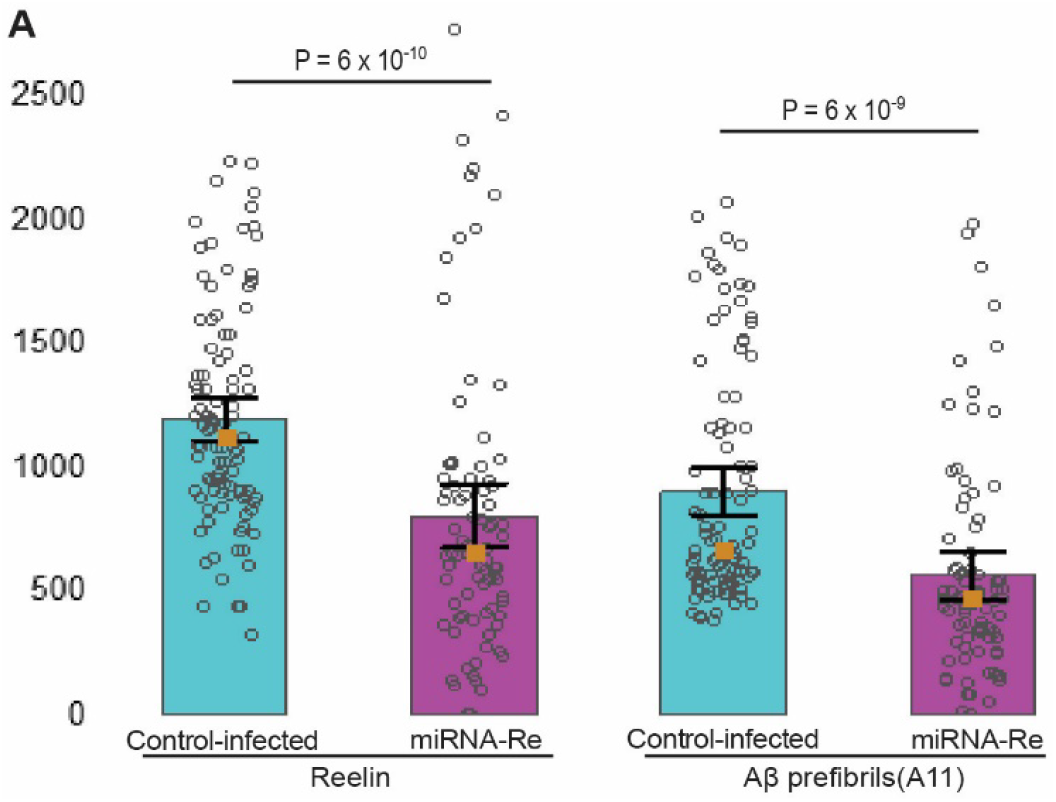
Lowering reelin in anterolateral entorhinal layer II-neurons by the experimental vector (miRNA-Re) leads to a 33 % reduction in levels of reelin, as measured by fluorescence-levels, in the hippocampal terminals of these neurons, relative to control. This results in a concomitant 37 % reduction of prefibrillar Aβ (A11). Means (µ) indicated by histogram, medians (Md) indicated by orange squares, 95% confidence interval for the means calculated from standard deviations (SD) are indicated by error bars. µ/Md/SD for levels of reelin: control vector-infected 1195,6 / 1122,37 / 439,24; miRNA vector-infected 801,99 / 654,08 / 587,98. µ/Md/SD for levels of prefibrillar Aβ A11: GFP 902,41 / 657,17 / 476,03; A11 miRNA 564,33 / 466,72 / 433,99.

## Supplementary tables

**Supplementary table 1.** Animals used with type of injection indicated. Age/time shown in days.

|  | Animal id | Sex | age at injection | time with virus | age at termination |
| --- | --- | --- | --- | --- | --- |
| miRNA-infected | 24868 | male | 36 | 78 | 114 |
|  | 24878 | Female | 39 | 75 | 114 |
|  | 26107 | Male | 37 | 60 | 97 |
|  | 26111 | Female | 42 | 55 | 97 |
|  | 26112 | Female | 42 | 55 | 97 |
|  | 26118 | Female | 43 | 54 | 97 |
|  | 26174 | Male | 35 | 59 | 94 |
|  | 26175 | Male | 35 | 59 | 94 |
|  | 26255 | Female | 24 | 58 | 82 |
| EGFP-infected | 26071 | Male | 24 | 61 | 85 |
|  | 26073 | Female | 25 | 60 | 85 |
|  | 26108 | Male | 38 | 59 | 97 |
|  | 26110 | Female | 42 | 55 | 97 |
|  | 26252 | Male | 23 | 59 | 82 |
|  | 26253 | Male | 23 | 59 | 82 |
|  | 26254 | Male | 23 | 59 | 82 |
|  | 24875 | Female | 37 | 77 | 114 |
|  | 25178 | Female | 36 | 56 | 92 |
|  | 25179 | Female | 35 | 57 | 92 |
| Double-infected | 25180 | Female | 33 | 59 | 92 |
|  | 25181 | Male | 33 | 59 | 92 |
|  | 25561 | Female | 32 | 63 | 95 |
|  | 25562 | Female | 33 | 62 | 95 |
|  | 24879 | Female | 38 | 76 | 114 |

**Supplementary table 2.**
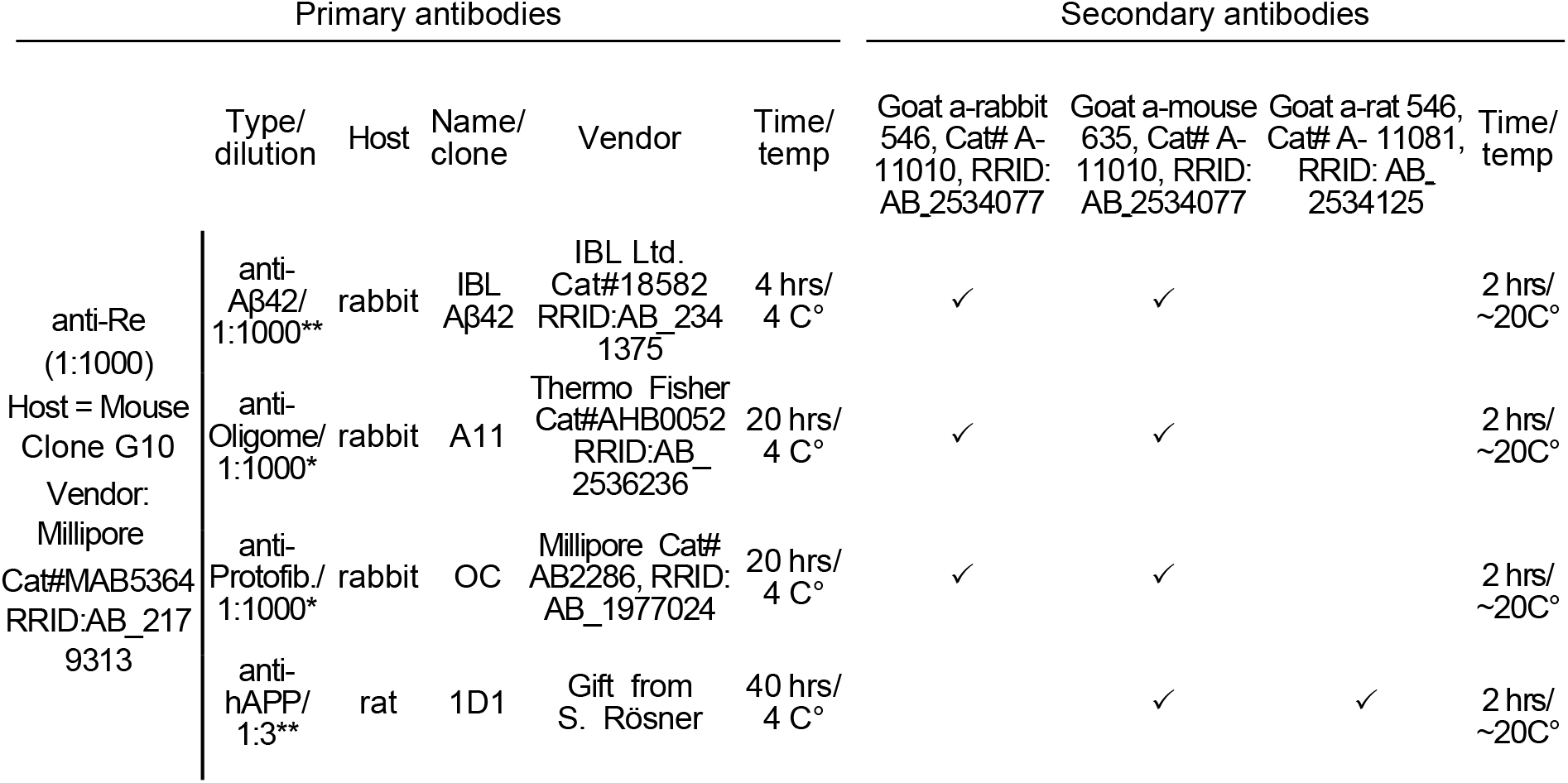
Overview of primary and secondary antibodies used. Note that we co-labeled with anti-reelin (leftmost column) for each of the other primary antibodies. Permeabilization agents used: *0.4% Saponin (VWR, Cat# 27534.187); **0.2% Triton-X100 (Millipore, Cat# 1.08603.1000). The anti-hAPP 1D1 antibody was a gift from Steffen Rossner at Paul Flechsig Institute for Brain Research, University of Leipzig, Germany(Höfling et al., 2016; PMID: 27470171); for this antibody we co-blocked (together with goat serum) with 3% donkey-Fab anti-rat IgG (Jackson ImmunoResearch Labs Cat# 712-007-003 RRIDAB_2340634).

**Supplementary table 3.**
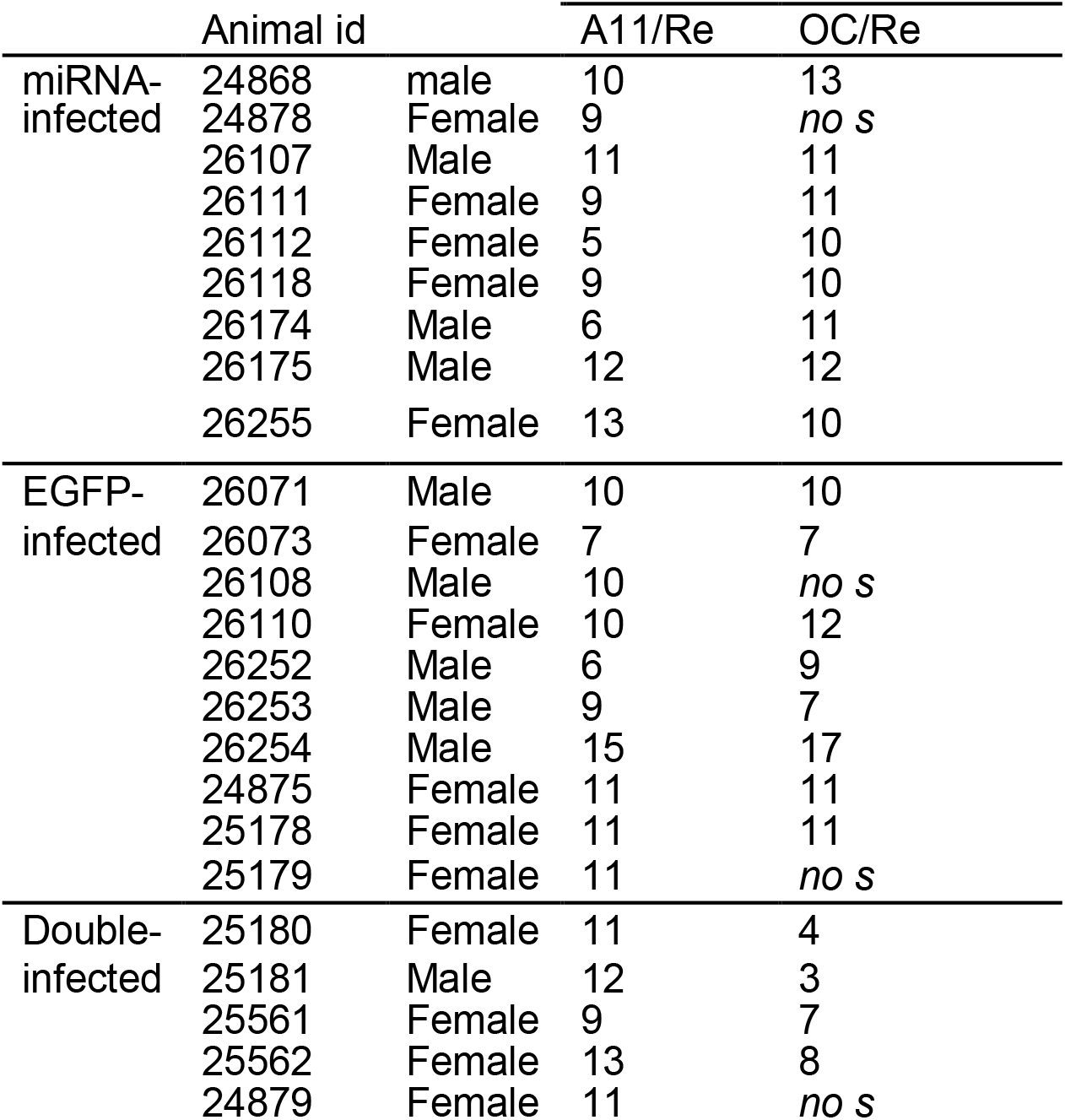
Number of sections used per animal to quantify levels of prefibrillar Aβ (A11), and protofibrillar Aβ (OC), both co-labeled with Reelin (Re), in ECLII projections in the outer molecular layer of the dentate gyrus (omlDG). For five IHC experiments the fluorescent signal was lost (*no s*) due to technical issues.

